# Repeat expansion and methylation state analysis with nanopore sequencing

**DOI:** 10.1101/480285

**Authors:** Pay Gießelmann, Björn Brändl, Etienne Raimondeau, Rebecca Bowen, Christian Rohrandt, Rashmi Tandon, Helene Kretzmer, Günter Assum, Christina Galonska, Reiner Siebert, Ole Ammerpohl, Andrew Heron, Susanne A. Schneider, Julia Ladewig, Philipp Koch, Bernhard M. Schuldt, James E. Graham, Alexander Meissner, Franz-Josef Müller

**Affiliations:** Department of Genome Regulation, Max Planck Institute for Molecular Genetics, (Berlin, Germany); Universitätsklinikum Schleswig-Holstein Campus Kiel, Zentrum für Integrative Psychiatrie gGmbH (Kiel, Germany); Oxford Nanopore Technologies (Oxford, UK); Kiel University of Applied Sciences, Institute for Communications Technology and Microelectronics (Kiel, Germany); Institute for Human Genetics, Ulm University and Ulm University Medical Center (Ulm, Germany); Department of Neurology, Ludwig-Maximilians-Universität (München, Germany); Central Institute of Mental Health, Medical Faculty Mannheim, Heidelberg University (Mannheim, Germany); HITBR Hector Institute for Translational Brain Research gGmbH (Heidelberg, Germany); German Cancer Research Center (DKFZ) (Heidelberg, Germany); Institute of Reconstructive Neurobiology, University of Bonn Medical Center (Bonn, Germany); Department of Stem Cell and Regenerative Biology, Harvard University (Cambridge, USA)

**Author notes:** Pay Gießelmann and Björn Brändl contributed equally to this work.

## Abstract

Expansions of short tandem repeats are genetic variants that have been implicated in neuropsychiatric and other disorders but their assessment remains challenging with current molecular methods. Here, we developed a Cas12a-based enrichment strategy for nanopore sequencing that, combined with a new algorithm for raw signal analysis, enables us to efficiently target, sequence and precisely quantify repeat numbers as well as their DNA methylation status. Taking advantage of these single molecule nanopore signals provides therefore unprecedented opportunities to study pathological repeat expansions.

---

The expansion of unstable genomic Short Tandem Repeats (STRs) causes more than 30 Mendelian human disorders^1^. For example, expansion of a GGGGCC-repeat [(G_4_C_2_)_n_] within the C9orf72 gene is the most frequent monogenic cause of Frontotemporal Dementia (FTD) and Amyotrophic Lateral Sclerosis (ALS; c9FTD/ALS; OMIM: # 105550)^2, 3^. Similarly, accumulation of a CGG motif in the FMR1 gene underlies the Fragile X Syndrome (FXS; OMIM # 300624), currently the most common identifiable genetic cause of mental retardation and autism^4^. In both prototypical repeat expansion disorders (Suppl. Discussion 1), recent evidence has suggested pronounced inter- and intraindividual repeat variability as well as changes in DNA methylation of the respective genomic regions to modulate disease phenotype^5^-^8^.

To overcome current difficulties in characterizing expanded STRs (Suppl. Discussion most notably we focused on three areas: i) optimization of Nanopore sequencing and signal processing to capture STRs ii) development and implementation of a target enrichment strategy to increase efficiency and iii) integration of expansion measurements with DNA methylation of the same molecule.

First, for benchmarking repeat expansion counting methods we constructed, verified and nanopore sequenced plasmids with several synthetic (G_4_C_2_)_n_-repeat lengths^9^. We analyzed our results with currently available STR quantification pipelines^10, 11^ but found those methods to become unreliable for more than 32 (G_4_C_2_)_n_-repeats with nanopore reads. To further improve the repeat analysis we developed a signal processing algorithm for a more exact quantification of STR numbers in raw nanopore signals (nanoSTRique: **nano**pore **S**hort **T**andem **R**epeat **i**dentification, **qu**antification &**e**valuation, Fig. 1a, Suppl. Fig. 1). Briefly (see Online Methods for details), reads spanning a STR location are identified by aligning the conventionally base-called sequences to a reference^12^. Next, nanoSTRique maps the upstream and downstream boundaries of each repeat more precisely with a signal alignment algorithm using SeqAn2^13^ and, as a third step, accurately quantifies the number of any given STR sequence with a Hidden Markov Model (Suppl. Fig. 1)^14^. Aggregated nanoSTRique repeat counts matched closely gel electrophoresis profiles from our synthetic repeats and could be confirmed on the single molecule level by manually counting repeat patterns in raw signal traces (Fig. 1d) (Suppl. Fig. 2, 3). Previously, repeat instability had been noted in Bacterial Artificial Chromosomes (BAC) containing expanded C9orf72 (G_4_C_2_)_n_-repeats (Online Methods)^15^. Analysing BAC clone 239 from a c9FTD/ALS patient (G_4_ C_2_)∼800 ^15^ with nanoSTRique we observed STR contractions in many reads and a secondary peak at 800 repeats (Fig 1c, Online Methods), while existing methods failed to mirror previously published Southern blot results (Suppl. Fig. 3).

**Figure 1.**
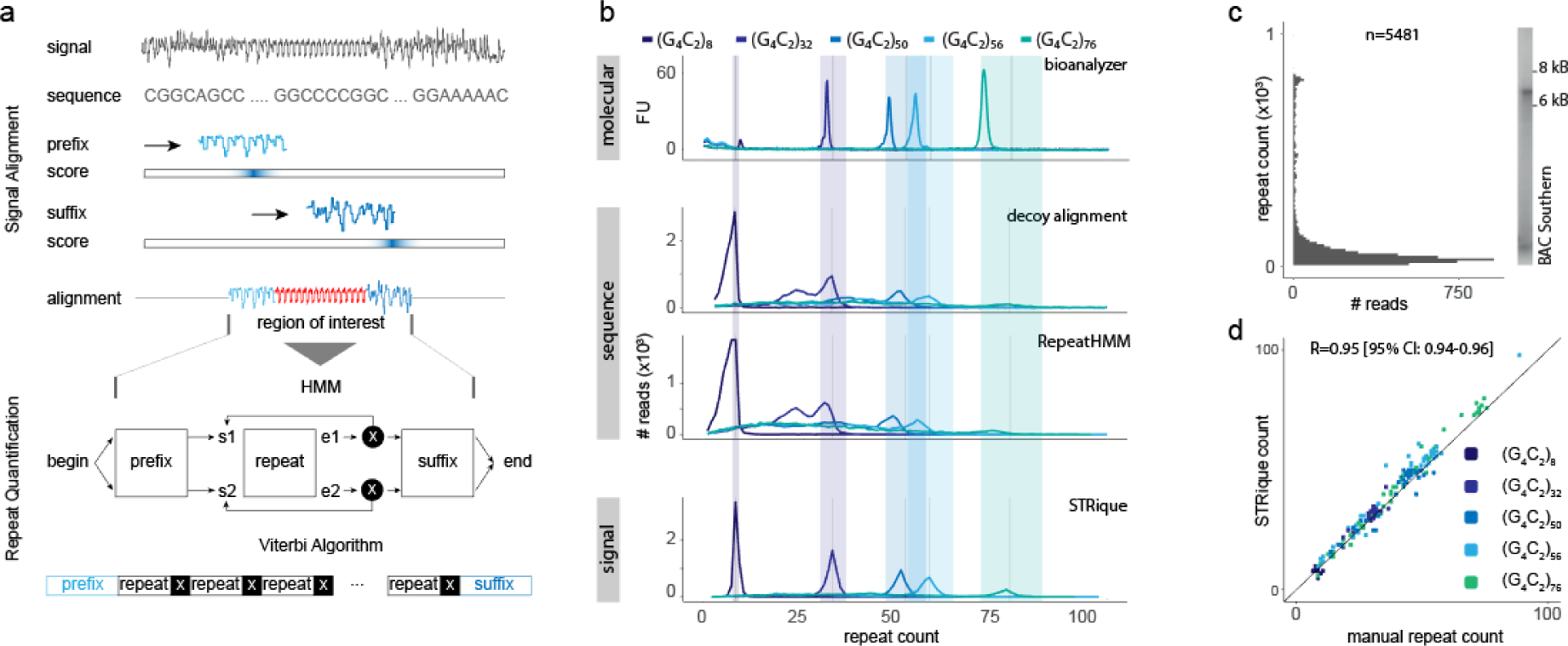
nanoSTRique: Generic repeat detection pipeline on raw nanopore signals. **a)**Repeat quantification by signal-alignment of flanking prefix and suffix regions and HMM based count on signal of interest. **b)** BioAnalyzer electropherogram, decoy alignment, RepeatHMM and nanoSTRique counts of synthetic (G_4_C_2_)_n_ repeats (10k random reads per barcode, +/- 10 % intervals around expected repeat length). **c)** Nanopore sequencing and analysis of BAC clone 239 from a c9ALS/FTD patient compared to cropped corresponding lane from Ref. 15 for illustration purpose. **d)** manual confirmation of detected repeat counts in synthetic repeats (n=16, 50, 49, 49, 47).

**Figure 2.**
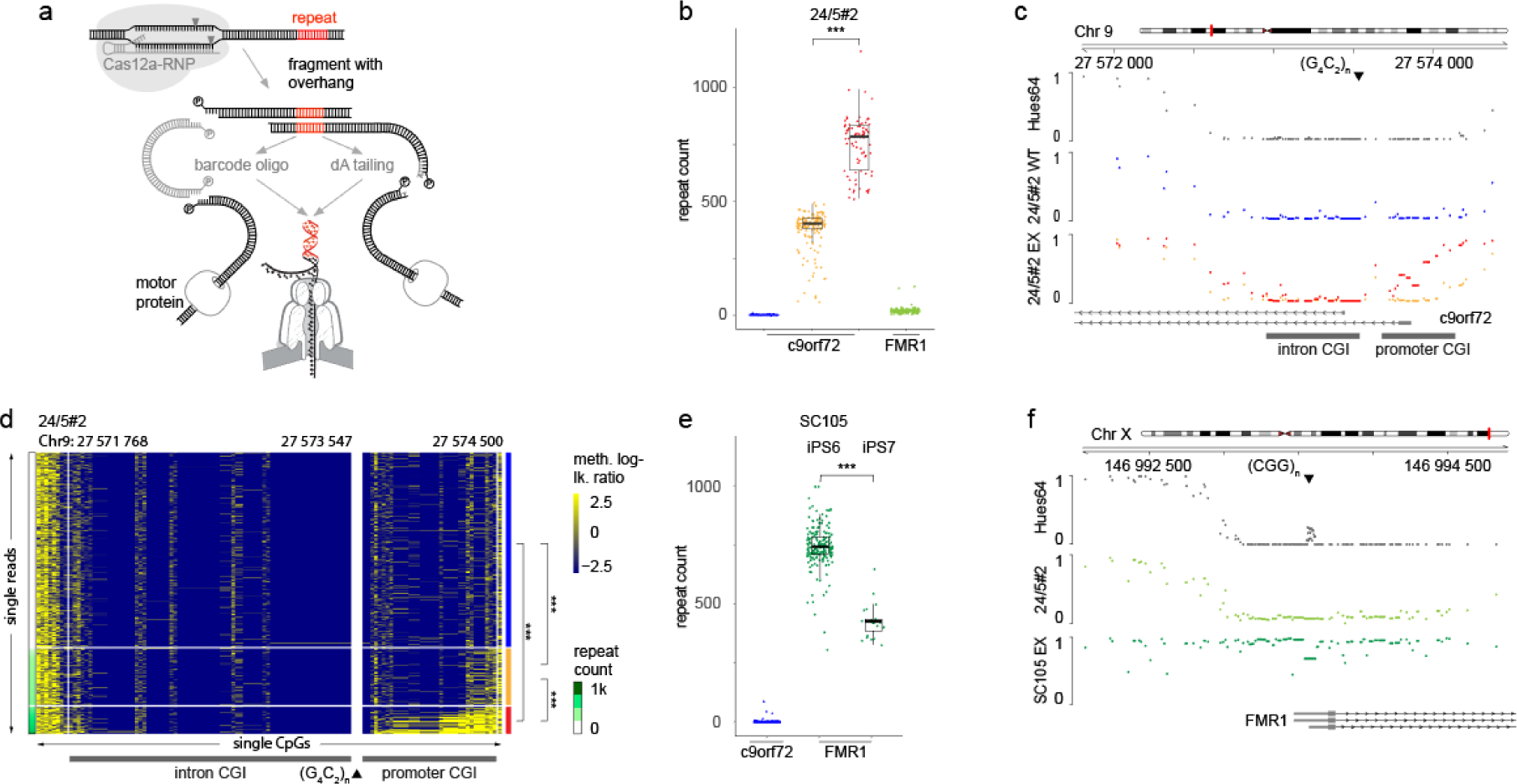
Targeted enrichment and nanopore sequencing with CRISPR-Cas12a. **a)** Illustration of the CRISPR-Cas12a target enrichment procedure **b)** Repeat quantification of sample 24/5#2 on c9orf72 and FMR1 loci revealing two distinct repeat bands of ∼450 and ∼750 (G_4_C_2_)_n_- repeats. (n=442,165,78,499; difference in repeat length 385 [95% CI: 361:404]) **c)** c9orf72 methylation status in Hues64 (WGBS), wild type allele and expanded allele of patient 24/5#2 (nanopore). **d)** Single read nanopore methylation of c9orf72 covering reads (n=769, row split 30,500) sorted by detected repeat length. (row:single read, column: single CpG, log-p > 2.5: methylated, log-p< −2.5: unmethylated, two sided wilcoxon rank sum test on mean promoter CGI methylation, median methylation difference [95% CI] wt-450: 6.9e-6 [3.5e-5 - 3.3e-2], wt-750: 0.33 [0.24 - 0.42], 450-750: 0.30 [0.20 - 0.42]) **e)** Repeat quantification of SC105iPS6/iPS7 sample on c9orf72 and FMR1 loci. (n=850,183,23; difference in repeat length −322 [95% CI: −351:-298]) **f)** FMR1 methylation status in Hues64 (WGBS), wild type allele of patient 24/5#2 and expanded alleles of samples SC105iPS6 and SC105iPS7^19^ (nanopore). (All box plots show median, box edges represent 1st and 3rd quartiles, whiskers extend to 1.5 x IQR; two sided wilcoxon rank sum tests, p-vals: * 0.05 - 0.01; ** 0.01 - 0.001; *** < 0.001)

Next we performed whole genome nanopore sequencing from c9FTD/ALS patient- derived DNA using four MinION flow cells yielding a total of 39 GBp (8M reads, 11 on target, none expanded). Our and similar nanopore whole genome sequencing results from others^16^ suggested a relevant bias against the detection of (G_4_C_2_)_n_- expansions with conventional molecular and bioinformatic nanopore workflows. To improve the coverage of any STR particularly the (G_4_C_2_)_n_-region in our proof of concept, we took advantage of the programmable CRISPR-Cas12a- ribonucleoprotein (Cas12a-RNP), which cleaves DNA via staggered double-strand breaks^17^. The Cas-system was applied to selectively target DNA sequences from a patient-derived induced pluripotent stem cell line (24/5#2) adjacent to the (G_4_C_2_)_n_- repeat resulting in unique 4 bp overhangs amenable to ligation of a linker oligo and subsequent attachment of the nanopore sequencing adapter (Fig. 2a, Online Methods). To further improve enrichment results we replaced the oligo adapter ligation step by adding Klenow fragment to fill in the Cas12a overhangs. The resulting dA-tailed DNA ends enabled even more efficient ligation of the sequencing adapters.

Additional dephosphorylation of all 5’ ends before Cas12a-RNP-digestion chemically protects DNA ‘background’ fragments from being ligated to sequencing adapters. Consequently, only those fragments cut by Cas12a-RNPs are capable of being sequenced by this procedure (Fig. 2a). Using this approach we were able to obtain a total of 1137 reads covering the (G_4_C_2_)_n_-repeat including 442 evaluable reads from expanded alleles (Suppl. Table 2, Suppl. Discussion 3). Consistent with Southern blot results from the same cell line (Suppl. Fig. 6 a-d), we found two distinct repeat expansion distributions (Fig. 2b). To explore the general applicability of our enrichment, sequencing and read processing we next tested two isogenic, patient-derived cell lines (SC105iPS6, SC105iPS7) carrying a FMR1-repeat expansion with an additional set of FMR1-targeting Cas12a-RNPs (Suppl. Table 1) and found two different repeat expansion distributions (Fig. 2e).

Lastly, epigenetic modification of both C9orf72 and FMR1 loci have been correlated with STR expansion status and patient characteristics in both disorders^7, 8^. We therefore combined single molecule CpG methylation analysis using nanopolish^18^ with our nanoSTRique results and found that in the 24/5#2 line all reads with STR expansions > 750 repeats showed significantly increased methylation at the promoter CpG island while all wild type reads and those with < 500 repeats were not or only partially methylated (Two sided wilcoxon rank sum test p < 0,001, Fig. 2C-D, Suppl. Fig. 4, Suppl. Discussion 4). Similarly, we found all FMR1 wild type alleles enriched from controls were unmethylated at a CpG island overlapping the CGG- STR. Consistent with previous findings^19^ and in our Southern blot analyses, all expanded alleles identified by nanoSTRique from both isogenic FXS-patient derived cell lines^19^, were determined to be methylated (Fig. 2f, Suppl. Fig. 6f-h).

Our results demonstrate the power of nanopore sequencing for the precise and multilayered molecular characterization of pathological short tandem repeat expansions. We have increased the enrichment for regions of interest on the background of the human genome approximately two to three orders of magnitude without any target amplification by using selective, multiplexed Cas12a-based chemical tagging of DNA fragments. Importantly, our method does not require any additional instruments in contrast to other previously reported enrichment strategies^20^ and enables reporting the DNA methylation status of the same alleles. The Cas12a-target enrichment and nanoSTRique can be rapidly adapted to any other genomic region of interest, ensuring broad applicability to overcome challenges associated with the single molecule analysis integrating genetic and epigenetic signals associated with unstable repeat expansions or any other as of yet ‘unsequenceable’ genomic regions in human health and disease. This type of analysis improves diagnostic workflows in regard to accuracy and resolution of unstable repeat expansion while enabling efforts to gain mechanistic insights into effects on differentiation, aging and future therapeutic agents that modify DNA methylation.

## 7. Acknowledgements

We are deeply thankful for the invaluable support by c9FTD/ALS and FXS patients and their families who donated fibroblast for this study. The C9orf72 BAC was generously provided by Robert Baloh and Shaughn Bell (Cedars Sinai Medical Center, Los Angeles, CA, USA). We are grateful to Jeanne Loring and Ai Zhang (The Scripps Research Institute, La Jolla, CA, USA) for providing us with hiPSC lines from a Fragile X patient (Supported by NIH R33MH087925-03). We thank Philip van Damme and Wim Robberecht (Laboratory for Neurobiology; VIB-KU Leuven Center for Brain & Disease Research, Leuven, Belgium) for providing the c9FTD/ALS patient-derived fibroblasts used for reprogramming. We acknowledge the expert assistance of the technical staff of the molecular genetics laboratory of the Institute of Human Genetics, Ulm. PK and JL acknowledge financial support by the Hector Stiftung II gGmbH. This work was supported by the Max Planck Society.

## 8. Author information

### Contributions

PG, BMS and FJM conceived the project. BB and RT performed cell culture, plasmid and BAC expansion and extraction. PG wrote the nanoSTRique pipeline. PG, BMS, CR, HK conducted additional bioinformatic analyses. PK and JL reprogrammed the c9FTD/ALS hiPSC from patient fibroblasts used in this study. ER, RB, AH, JEG developed the Cas12a-RNP protocol. BB further developed the Cas12a protocol with c9FTD/ALS and FXS patient-derived DNA and performed all nanopore library preparation and nanopore sequencing for the results presented in this manuscript, RT and CG worked on optimization of aspects of the enrichment protocol. GA and RS conducted diagnostic testing of the repeat expansions by Southern Blot and PCR analyses, SS, RS, OA and GA provided clinical and diagnostic advice. PG, BMS, AM and FJM wrote the manuscript. FJM oversaw the study. All authors contributed to the editing and completion of the manuscript.

## 9. Data availability

All sequencing data points generated in this study (i.e. nanopore reads, raw and base called) and utilized for the determination of FMR1 and C9orf72 repeat expansion lengths and methylation status will be made public upon acceptance through a public repository (e.g. EGA). Whole genome sequencing data generated for this study will be made accessible to researchers upon request under a Data Access Agreement similar to the procedure required for managed access by the European Genome-phenome Archive. (for details see: https://www.ebi.ac.uk/ega/submission/data_access_committee/policy_documentation).

## References

1. Gatchel, J. R. & Zoghbi, H. Y. Diseases of unstable repeat expansion: mechanisms and common principles. Nat Rev Genet 6, 743–755 (2005).

2. Renton, A. E. et al. A hexanucleotide repeat expansion in C9ORF72 is the cause of chromosome 9p21-linked ALS-FTD. Neuron 72, 257–268 (2011).

3. DeJesus-Hernandez, M. et al. Expanded GGGGCC hexanucleotide repeat in noncoding region of C9ORF72 causes chromosome 9p-linked FTD and ALS. Neuron 72, 245–256 (2011).

4. Verkerk, A. J. et al. Identification of a gene (FMR-1) containing a CGG repeat coincident with a breakpoint cluster region exhibiting length variation in fragile X syndrome. Cell 65, 905–914 (1991).

5. van Blitterswijk, M. et al. Association between repeat sizes and clinical and pathological characteristics in carriers of C9ORF72 repeat expansions (Xpansize-72): a cross-sectional cohort study. Lancet Neurol 12, 978–988 (2013).

6. Xi, Z. et al. Hypermethylation of the CpG Island Near the G4C2 Repeat in ALS with a C9orf72 Expansion. The American Journal of Human Genetics 92, 981–989 (2013).

7. Russ, J. et al. Hypermethylation of repeat expanded C9orf72 is a clinical and molecular disease modifier. Acta Neuropathol. 129, 39–52 (2015).

8. Hornstra, L. K., Nelson, D. L., Warren, S. T. & Yang, T. P. High resolution methylation analysis of the FMR1 gene trinucleotide repeat region in fragile X syndrome. Hum Mol Genet 2, 1659–1665 (1993).

9. Mizielinska, S. et al. C9orf72 repeat expansions cause neurodegeneration in Drosophila through arginine-rich proteins. Science 345, 1192–1194 (2014).

10. Liu, Q., Zhang, P., Wang, D., Gu, W. & Wang, K. Interrogating the ‘unsequenceable’ genomic trinucleotide repeat disorders by long-read sequencing. Genome Medicine 2017 9:19, 65 (2017).

11. Dashnow, H. et al. STRetch: detecting and discovering pathogenic short tandem repeat expansions. Genome Biol 19, 121 (2018).

12. Li, H. Minimap2: pairwise alignment for nucleotide sequences. Bioinformatics 3, 321 (2018).

13. Reinert, K. et al. The SeqAn C++ template library for efficient sequence analysis: A resource for programmers. J. Biotechnol.261, 157–168 (2017).

14. Schreiber, J. & Karplus, K. Analysis of nanopore data using hidden Markov models. Bioinformatics 31, 1897–1903 (2015).

15. O’Rourke, J. G. et al. C9orf72 BAC Transgenic Mice Display Typical Pathologic Features of ALS/FTD. Neuron 88, 892–901 (2015).

16. Ebbert, M. T. W. et al. Long-read sequencing across the C9orf72 ‘GGGGCC’ repeat expansion: implications for clinical use and genetic discovery efforts in human disease. Mol Neurodegeneration 13, 46 (2018).

17. Zetsche, B. et al. Cpf1 Is a Single RNA-Guided Endonuclease of a Class 2 CRISPR-Cas System. Cell 163, 759–771 (2015).

18. Simpson, J. T. et al. Detecting DNA cytosine methylation using nanopore sequencing. Nat Methods (2017). doi:10.1038/nmeth.4184

19. Boland, M. J. et al. Molecular analyses of neurogenic defects in a human pluripotent stem cell model of fragile X syndrome. Brain 140, 582–598 (2017).

20. Gabrieli, T. et al. Selective nanopore sequencing of human BRCA1 by Cas9-assisted targeting of chromosome segments (CATCH). Nucl. Acids Res. 46, e87–e87 (2018).

