## Supplementary material for "Repeat expansion and methylation state analysis with nanopore sequencing"

* corresponding author

### Raw nanopore signal processing

We propose a raw nanopore signal based pipeline to detect arbitrary repeat patterns based on signal alignment and HMM driven quantification. We adapted the SeqAn2 C++ library (https://github.com/seqan/seqan)[1] to support semi global alignment of generic signals using the score function:

$$s_{i,j} = max \begin{cases} c - |x_i - y_j| \\ -c \end{cases}$$

The distance score in the dynamic programming matrix of two signal values $x_i$, $y_i$ is computed as their absolute difference subtracted from a constant offset to map more similar values to a positive score and less similar values to negative ones. The negative score is capped at a constant threshold. In contrast to the regular dynamic time warping (DTW) algorithm we applied affine gap costs without penalizing the first and last gap of the alignment (semi-global), allowing us to search for patterns of interest in long nanopore raw signals (Fig. 1a, Suppl. Fig. 1a-b).

Given the prior knowledge of the prefix and suffix sequence of the targeted repeat we simulate these sequences according to the pore model and use the signal alignment to map prefix and suffix in the signal space. A compound profile HMM of prefix, repeat and suffix signal states quantifies the repeat by counting the iterations in the repeat part of the model after computing the viterbi path of the entire model given the observed signal (Suppl. Fig. 1a).

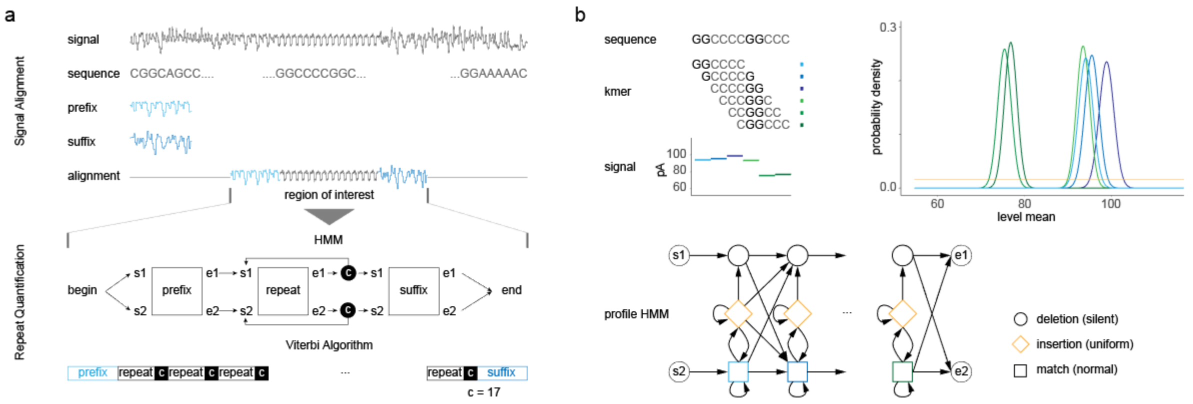


**Supplementary Figure 1:** Nanopore signal processing with nanoSTRique

a) Signal alignment detecting prefix and suffix flanking the repeat using a capped distance score function and a semi-global alignment. A compound profile HMM of prefix, a single repeat and the suffix sequence assigns either prefix, repeat or suffix label to each signal value. Repeat counts are obtained through dummy states between repeat and suffix. b) Nanopore signal profile HMM with normal distributed match states and uniform distributed insertion states.

The compound HMM is build of generic profile HMM blocks similar to the architecture proposed in[2]. For a given profile sequence the expected nanopore signal values are extracted from a pore model. The emission probabilities of the match states are then expressed by normal distributions around these values. Insertion states with an uniform distribution between model minimum and maximum allow to compensate for intermediate state and noise measurements. Silent deletion states enable the model to dodge states without observations. Beside the repeat count nanoSTRique provides the alignment scores of prefix and suffix signal. In this study we used a thresholds of 3.8 for human and 4.0 for plasmid and BAC samples to discard low quality counts.

#### MinION base-calling and alignment

All MinION runs were base-called using ONT Albacore 2.3.3 with disabled quality filtering. Alignments were made using minimap2 (v2.12) with default parameters to identify reads spanning the C9orf72 and FMR1 regions of interest and determine the read strand.

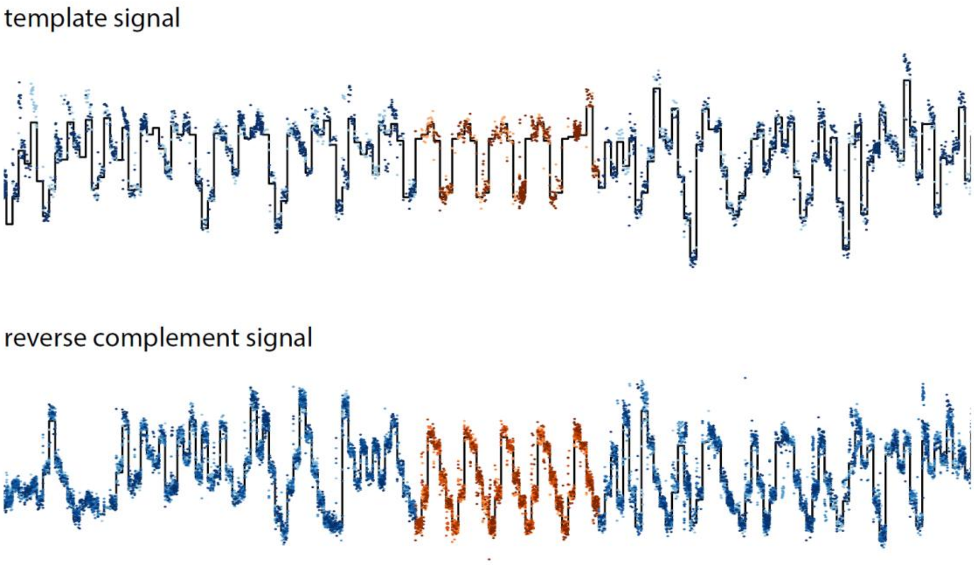


**Supplementary Figure 2:** Nanopore raw signal at the C9orf72-$(G_4C_2)_n$-locus in NA12878 cells

Compound multi signal HMM alignment of publicly available[3] raw traces from two template and eight complement reads from the NA12878 cell line shows matching signal pattern in all reads. Displayed are the current measurements as dots and the model signal as black line. Blue dots indicate current measurements identifies as prefix or suffix (see Suppl. Fig.1). Red dots indicate raw current measurements identified by nanoSTRique as belonging to the C9orf72-$(G_4C_2)_n$-STR. NanoSTRique detects in this case five $(G_4C_2)$-repeats in the reads.

#### Repeat simulation with Nanopore SimulatION

For development and in silico experimental setup validation repeats were simulated with our Nanopore SimulatION tool (https://github.com/crohrandt/nanopore_simulation). It takes a reference genome fasta file, and a configuration (.sic file) from a previous real nanopore experiment as input for a realistic raw signal simulation of nanopore sequencing data. The configuration may be modified within an ini file at simulation time. The parameters include read length distribution, signal capture characteristics and experiment metadata. Furthermore the ini file is a mean of controlling simulation parameters as the induced signal error, the simulation of only specific sequences in full length or embedding a ground truth sequence into the output. For simulation metrics a pre derived model of all 6-meres is used. Nanopore SimulatION provides fast5 files that are compatible to the standard software pipeline used for basecalling.

#### RepeatHMM repeat detection

RepeatHMM[4] was used as first instance to quantify our synthetic $(G_4C_2)_n$-repeat plasmid data. The software, initially validated with data only from trinucleotide (SCA3/ATXN3) and pentanucleotide (SCA10/ATXN10) tandem repeat expansions, was extended by user defined repeat parameters in this study. Using this feature it was possible to detect the $G_4C_2$-hexanucleotide. Furthermore the software was modified to also output a per read repeat count as it originally only outputs a distribution over the whole amount of reads passed. The modified version is available at https://github.com/giesselmann/RepeatHMM.

#### Alignment based repeat detection

We next implemented a naïve repeat detection approach by base calling reads and aligning against a set of ‘decoy’ references with each possible repeat length inspired by the STRetch method proposed by Dashnow and colleagues[5]. The repeat count is obtained from the best matching reference without allowing multiple alignments.

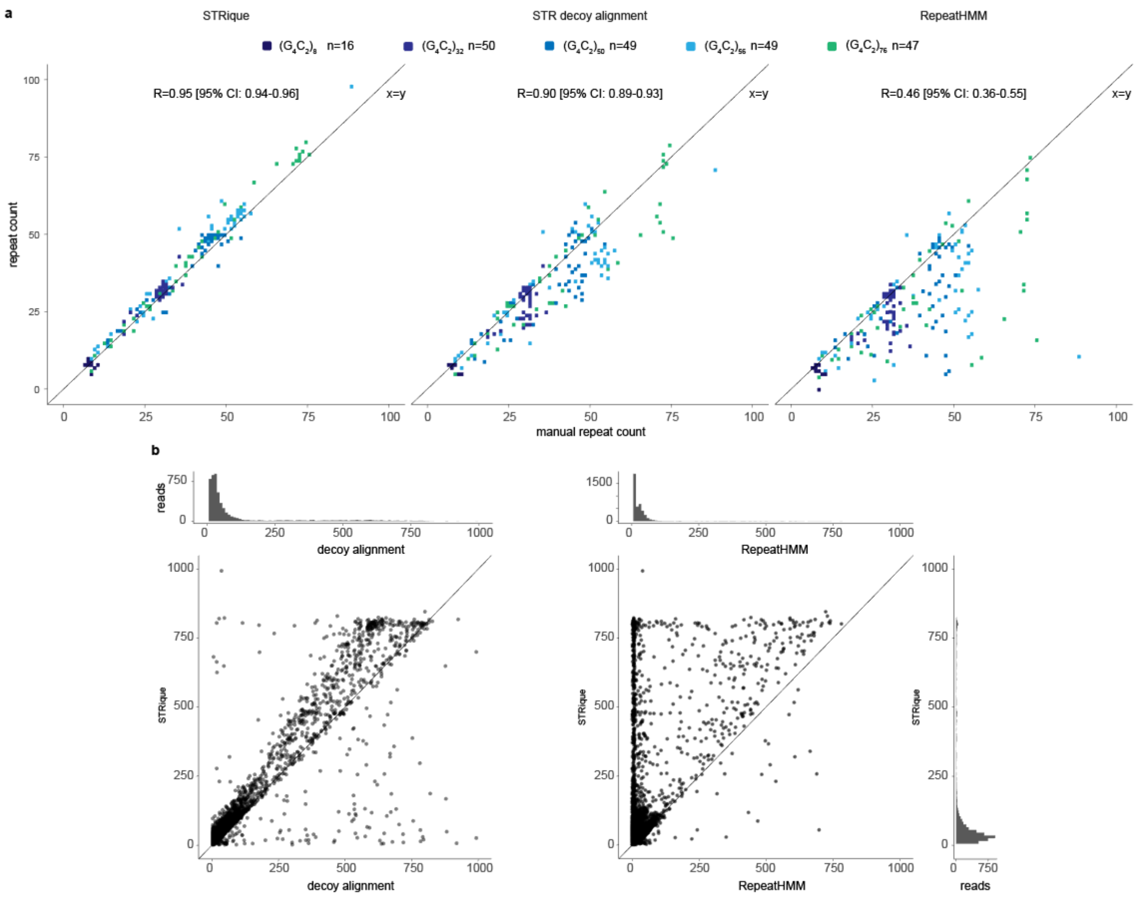


**Supplementary Figure 3** Evaluation of repeat quantification approaches

**a)** Manual counted set of 50 plasmid reads per repeat length in (8,32,50,56,76) on x-axis correlating with STRique raw signal pipeline, albacore base calling and decoy alignment approach and RepeatHMM sequence pipeline on y-axis. Only data points shown which could be evaluated with all four methods. Less counts for 8 repeats due to the limited visibility in the raw signal. Sequence based methods show an overall trend towards underestimation of the repeat length. **b)** Correlation of STR decoy alignment and RepeatHMM repeat counts with STRique pipeline on BAC data. These results extend the observations made with the synthetic repeats for repeat expansion numbers greater than 76 in. n=5481 reads passing all three pipelines.

We note, that RepeatHMM was only developed and validated for trinucleotide (SCA3/ATXN3) and pentanucleotide (SCA10/ATXN10) repeat expansions, yet its parameter and output set could be adapted to also accommodate hexanucleotide repeat expansions. As with the synthetic repeat sequences, STR decoy alignment assigns in general lower repeat counts to most sequences in comparison to nanoSTRique with the divergence becoming more pronounced after repeat numbers increase from ~250 to up to 800. Interestingly, a subgroup of repeat expansion counts remain in agreement between STR decoy alignment and nanoSTRique.

Notably only nanoSTRique replicates the peak around ~800 repeats that mirrors the expected repeat expansion maximum from Southern blots from the c9FTD/ALS patient and the BAC derived from the patient which we used for this study (see for comparison Fig. 1C Patient 29 in Sareen et al. 2013[6] and Fig. 1S lane 24 in O'Rourke et al 2015[7])

#### Methylation detection on expanded reads

Methylation tracks were generated by masking the repeat signal section of expanded reads in the raw fast5 (script fast5Masker.py), re-basecalling and alignment with graphmap (v0.5.2) and finally nanopolish (v0.9.0) methylation calling each with default parameters. The raw nanopore methylation log-likelihood ratio per CpG is interpreted as methylated for values greater 2.5 and unmethylated for values less than -2.5. Intermediate values are used in the single read heat maps but discarded in the region methylation tracks.

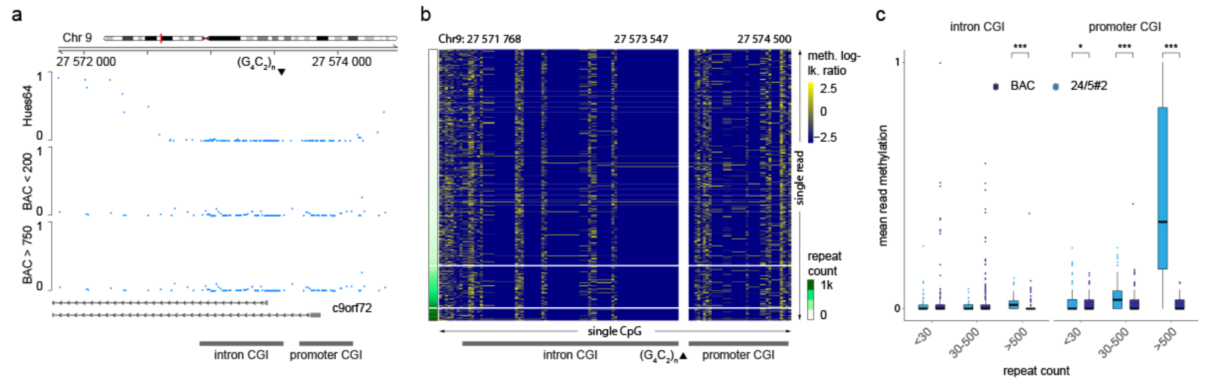


**Supplementary Figure 4** Nanopore single read methylation in BAC data

**a)** Methylation status of c9orf72 region in BAC data for repeats < 200 and > 750 and Hues64 (WGBS) control **b)** single read methylation on a random sample of 500 reads sorted by repeat count showing no methylation signal correlating with the repeat count (row split 200 and 750 repeats). **c)** mean methylation per read binned by detected repeat length (24/5#2 n = 793 0-30; 235 30-500; 106 >500 BAC n = 1095 <*30*; 2223 *30-500*; 347 >*500*) of intron and promoter CGI for BAC and patient 24/5#2 (two sided wilcoxon rank sum test, p-val: * 0.05 - 0.01; ** 0.01 - 0.001; *** < 0.001; box plots show median, box edges represent 1st and 3rd quartiles, whiskers extend to 1.5 x IQR; median methylation difference intron (95%CI): *>500* -0.2e-4 [CI: -1.5e-2:-4.2e-5] promoter: *<30* -3.8e-5 [CI: -6.5e-5:-1.2e-5] *30-500* -1.3e-4 [CI: -3.4e-2:-7.4e-5] *>500* -0.3 [CI: -0.4:-0.2]).

Effect sizes of mean read methylation levels are given as median of differences using standard R wilcox.test with conf.int set to true.

#### Preparation of high molecular weight DNA from cultured c9FTD/ALS patients derived cells

High Molecular Weight DNA (HMW DNA) was prepared with a modified phenol-chloroform extraction method [8]. Briefly $2x10^7$ cells (undifferentiated hiPSC) were detached with TrypLE Select (Thermo Fisher Scientific) for 3 min at 37 °C. Enzymatic reaction was stopped with DMEM/F-12 (Thermo Fisher Scientific) and the cell suspension was centrifuged for 3 min at 260 x g. Supernatant was discarded and cells were resuspended in 100 µl of 1X PBS. Cells were lysed by adding 10 mL of TLB solution composed of 10 mM Tris-Cl pH 8, 25 mM EDTA pH 8, 0,5 % SDS (w/v) and 20 µg/mL RNase A (Qiagen) for 1 h at 37 °C. Proteins were subsequently digested at 50 °C for 3 h using 50 µl Proteinase K (> 600 mAU/mL, Qiagen). The viscous solution was transferred into a 50 mL falcon tube containing 5 g of phase lock gel (High Vacuum Grease, Dow Corning) and 10 mL of ultra pure saturated phenol (Thermo Fisher Scientific) was added. Samples were placed onto a rotator at 40 rpm for 10 min until a fine emulsion had formed and a phase separation was performed by centrifugation at 2.800 x g for 10 min. The aqueous phase was carefully poured into a fresh 50 mL Falcon tube containing 5 g of phase lock gel followed by a second phase separation using 5 mL of ultra pure saturated phenol and 5 mL chloroform (Merck). Samples were mixed and centrifuged as described above. The aqueous phase was poured into a fresh 50 mL falcon tube and the genomic DNA was precipitated using 2.7 mL of 7.5 M ammonium acetate and 30 mL of ice-cold ethanol (absolute) with subsequent gentle inversion for 10 times. Precipitated DNA was spooled out of solution using a glass rod and carefully submerged in 80 % ethanol. HMW DNA were transferred into a 1,5 mL DNA LoBind tube (Eppendorf) containing 1 mL of 80 % ethanol and centrifuged at 16.000 x g for 10 min. Supernatant was removed and the DNA pellet was dried at 40 °C for 5-10 min. Rehydration of DNA was done at 50 °C for 1-2 h using 100 µl of 10 mM Tris-HCl pH 8. Samples were stored at 4 °C until use.

#### Selective ligation of genomic fragments with CRISPR-Cas12a-RNPs mediated DNA cleavage.

We designed CRISPR-Cas12a crRNAs targeting genomic regions adjacent to the C9orf72 $(G_4C_2)_n$- and FMR1 CGG-repeats with the ChopChop online tool (http://chopchop.cbu.uib.no, Suppl. Table 1), [9,10]. creating staggered 4 bp overhangs at the 5' end. We note that Cas12a had been introduced into the literature as Cpf1 (**C**RISPR from **P**revotella and **F**ranciscella**1**[11]) but is now more frequently referred to as Cas12a[12,13]. Next we designed unique ssDNA oligomere ("bottom strand", IDT Technologies) in parts complementary to a universal ssDNA barcode (NB01) referred as "top strand" based on the ONT Native Barcoding Kit 1D (EXP-NBD103). Oligomere were resuspended in 10 mM nuclease-free Tris-Cl pH 8 to a final concentration of 100 µM and subsequently used for barcode annealing. Briefly 44 µl of top strand were combined with 40 µl of the respective bottom strand and 16 µl of nuclease-free duplex annealing buffer (IDT). Samples were heated for 2 min at 95 °C in a thermocycler. Mixture was cooled down to 25 °C over 70 min with a cooling rate of approximately -1 °C per minute. Successfully annealed barcodes serve as a linker that provide complementary overhangs to the respective Cas12a cut site and to the BAM 1D adapter thus allowing a sticky-end ligation.

Preparation of programmed Cas12a nucleoprotein complex (Cas12a-RNP) was performed by combining the EnGen Lba Cas12a (NEB) with IDT Alt-R™ Cpf1 crRNA that consists of a target specific 21 bp protospacer domain and a constant 20 bp loop domain. Briefly 2 nmol of lyophilized crRNA were reconstituted with 20 µl of 10 mM nuclease-free TE buffer pH 7.5 to obtain a 100 µM solution. Next, crRNA were pooled and adjusted to 10 µM using nuclease-free water. Formation of secondary RNA structure were done in 1X NEB CutSmart buffer with a final crRNA concentration of 500 nM. The Sample was heated for 6 min at 90 °C using a thermomixer and snap-cooled on wet ice before adding 0,25 µl of Cas12a enzyme (100 µM stock) to obtain a final concentration of 500 nM. Cas12a-RNPs were formed for 20 min at room temperature (RT) and assembled complex was stored on ice until use.

HMW DNA were digested with 10 µl of FastDigest BamHI (Thermo Fisher Scientific) in 1X NEB CutSmart Buffer for 12 h at 37 °C and subsequently heat inactivated at 80 °C for 5 min. As a next step the 5' ends were dephosphorylated for 30 min at 37 °C using 4 µl of quick calf-intestinal alkaline phosphatase (CIP, NEB) followed by a heat inactivation step for 5 min at 80 °C. Sample were equilibrated to room temperature prior Cas12a-RNP incubation. To allow sufficient Cas12a cleavage, dephosphorylated DNA were supplemented with 10 µl of Cas12a-RNPs and incubated for 1 h at 37 °C. Cleavage reaction was continued at 4 °C overnight (ON).

Next day the cleaved DNA sample was carefully cleaned and concentrated by using 1X volume of Agencourt AMPure XP beads (Beckman Coulter). DNA binding to AMPure beads were done for 10 min at RT followed by quick pulse centrifugation. Next, beads were immobilized at a DynaMag Spin magnet (Thermo Fisher Scientific) for 5 min, and the sample were washed with 80 % ethanol according to the manufacturer's protocol. Residual ethanol was completely removed by pipetting and beads were air dried for 60 sec. AMPure beads were carefully resuspended in 40 µl of pre-warmed (37 °C) 0,1X TE buffer using large bore tips, followed by a 10 min incubation step at 37 °C. Sample tube were placed on the magnet and eluted DNA were used for barcode adapter ligation.

Annealed barcode adapter (see above) were pooled and adjusted to 1 µM working concentration. A total of 0,2 µl pooled barcode mix were added to the cleaved DNA. Sample were incubated at 65 °C for 5 min in a thermocycler and equilibrated to RT before adding 60 µl of NEB Blunt/TA Ligase Master Mix (NEB) followed by an additional incubation for 30 min at RT. Next, 20 µl of BAM 1D sequencing adapter from the ONT Native Barcoding Kit 1D (EXP-NBD103) were added to the above mix and subsequently ligated at RT for 30 min.

The sequencing library were purified again by adding 0.4X volume of Agencourt AMPure XP beads for 10 min at RT. Excess of BAM 1D were washed away with adapter bead binding (ABB) solution according to the manufacturer's protocol. Final sequencing library were eluted from the magnetic beads using 18,5 µl of prewarmed elution buffer (ELB) from the ONT Ligation Sequencing Kit 1D (SQK-LSK108) at 37 °C for 20 min. The sample tube were placed on the magnet and eluted library were transferred to a fresh DNA LoBind tube at 4 °C. Sample was then quantified using a

Qubit fluorometer together with the Qubit dsDNA BR Assay Kit (both Thermo Fisher Scientific) and a total of 17,5 µl of library were combined with 35 µl of running buffer (RBF) and 22,5 µl of library loading beads (LLB) from the EXP-LLB001 kit. After removal of the AMPure beads, a ONT SpotON Flow Cell R9.4(FLO-MIN106) was primed and loaded following the manufacturer's instructions with no modifications to the procedure. Suppl. Table 2 gives an overview of the flow cells, DNA samples, yield and run characteristics used for this study.

Improved enrichment experiments were based on a modified Cas12a protocol that contained a dA-Tailing reaction, yield substantially higher amount of on target reads and took advantage of an improved sequencing kit (SQK-LSK109). Initial BamHI digestion, the use of barcode adapter and ON incubation with RNP was no longer required. Instead HMW DNA was diluted to a final concentration of 200 ng/µl and a total of 5µg DNA were dephosporylated with 2 µl Quick CIP in 1X NEB CutSmart buffer with same parameters as described above. Assembled RNP complex were added to the dephosphorylated DNA together with 1 µl of 1 mM dNTP mix and 0,5 µl of 10 mM dATP and reaction volume were adjusted to 49 µl using nuclease-free water. Briefly 5 U of NEBs Klenow Fragment (3'→5' exo-) were added to the sample and both dA-tailing as well as RNP cleavage were done in parallel while incubating the sample at 37 °C for 15 min followed by a second incubation at 65 °C for 7 min. Instead of using the BAM 1D adapter the dA-tailed library were incubated with an adapter ligation mix composed of 25 µl ligation buffer (LNB), 10 µl Quick T4 DNA Ligase (NEB), 10 µl nuclease-free water and 5 µl of adapter mix (AMX). Adapter ligation was done for 20 min at RT and mixture was subsequently diluted with 1X volume of 10 mM nuclease-free Tris-Cl pH 8. Following ligation sample were incubated with 0,3X volume of AMPure beads for 10 min at RT and washed twice with 250 µl long fragment buffer (LFB) following the manufacturer's instructions. Washed library were eluted with 16 µl elution buffer (EB) and 1 µl was used for Qubit quantification as described previously. Priming of the ONT SpotON Flow Cell R9.4(FLO-MIN106) were performed according on the manufacturer's protocol with minor modification. Prior sample loading a SpotON priming mix composed of 20 µl sequencing buffer (SQB), 0,4 µl sequencing tether (SQT) and 19,6 µl nuclease-free water were added dropwise to the SpotON port. The eluted DNA library were supplemented with 25 µl SQB together with 10 µl loading beads (LB) and immediately loaded as described earlier.

#### Nanopore whole genome, plasmid and BAC sequencing

For whole genome sequencing, we generated sequencing libraries from BamHI digested genomic DNA with the ONT Ligation Sequencing kits 1D (SQK-LSK108) following the manufacturer's instructions without modifications.

For sequencing our synthetic $(G_4C_2)_n$ repeats containing plasmids, we first linearized the plasmids with NdeI (pcDNA3.1(+)) or ScaI (pCR Script Amp-BE), heat inactivated the restriction enzymes and cleaned-up the samples using the NucleoSpin Gel and PCR clean-up following the manufacturer's instructions. Linearized plasmid DNA, was further processed with the ONT Ligation Sequencing kits 1D (SQK-LSK108) following the the manufacturer's instructions without modifications. In some instances, the plasmids were barcoded using adapters included in the 1D native barcoding kit (EXP-NBD103) following the manufacturer's instructions without modifications before the 1D ligation sequencing kit was applied.

DNA obtained from pCC1-BAC clone 239 (O'Rourke 2015) was sequenced using the ONT 1D rapid sequencing kit (SQK-RAD002) following the manufacturer's instructions without modifications. All samples were supplemented with beads from the library loading bead kit (EXP-LLB001) prior sample loading on the flow cells.

#### Synthetic $(G_4C_2)_n$-repeat cloning, modification and visualisation

For cloning $(G_4C_2)_n$ fragments carrying a defined repeat number we used recursive directional ligation as described in Mizielinska et al. 2014 [14]. Briefly, complementary DNA oligonucleotides  (Suppl. Table 3) containing three or four $G_4C_2$ -repeats flanked 5' by BamHI and BspQI (GCTCTTCC*GGCC) recognition sites and 3' by EcoO109I (GG*GGCCT) and NotI recognition sites, were annealed and ligated into the vector pCR Script Amp-BE using the BamHI and NotI sites. Inserts were excised by digestion of the vector with BspQI (LguI) and EcoO109I (Thermo Fisher Scientific), electrophoretically separated, purified from agarose gels (Qiagen

MinElute Gel Extraction Kit), subsequently religated into the BspQI(LguI)-linearized pCR Script Amp-BE vector. Six cycles of recursive directional ligation gave rise to increasing insert sizes of up to 100 repeat elements. Transformations were performed using recombination-deficient Stbl3 E. coli (Thermo Fisher Scientific) at 30 °C for longer repeats to minimize retraction of repeats. DNA was extracted using Plasmid Mini/Maxi kits (QIAGEN), following the manufacturer's instructions. Constructs were screened using standard restriction enzyme digest and agarose gel electrophoresis (2 % or 4 %). Next we sequence-verified (GENterprise Genomics, Mainz, Germany) plasmids with up to 32 repeats, to confirm repeat size and lack of interruptions (data not shown). In case of the 7, 8, 32, 76 and 100 repeats, the appropriate fragment length was excised from pCR Script Amp-BE and religated into pcDNA3.1(+) via BamHI/NotI. For exact repeat length visualisation and comparison with nanopore results from the same plasmids, we followed procedures outlined by Kwok and colleagues for the detection of the FMR1 repeat expansion[15]with one relevant modification. Instead of using PCR-amplified fragments from the repeat region, we used fragments cut out of our synthetic pCR Script Amp-BE $(G_4C_2)_n$-plasmids with restriction enzymes. We analysed repeat inserts from different repeat lengths (8, 32, 50, 56, 76), which were excised with BamHI/NotI or NdeI/ScaI from the pCR Script Amp-BE plasmids and analyzed with a 2100 Agilent Bioanalyzer on a 1000 DNA gel cassette following the manufacturer's instructions (Agilent, Böblingen, Germany). Bioanalyzer raw data was normalized and plotted using an in-house script following Agilent's analysis steps (Suppl. data).

#### C9orf72 BAC expansion and DNA extraction

A 174 kb bacterial artificial chromosome (pCC1-BAC clone 239) containing a $(G_4C_2)$-repeat expansion from a c9FTD/ALS patient [7] was amplified as previously described. Briefly, transfected DH10B T1 cells (Thermo Fisher Scientific) were grown on agar plates (LB broth with agar) and in LB broth containing 12.5 ng/µl Chloramphenicol at temperatures < 30 °C as higher temperatures lead to repeat contraction and/or loss of the BAC. Extraction of BAC DNA was done using the Qiagen Large-Construct Kit (Qiagen) including an ATP-dependent Exonuclease step for sufficient removal of genomic DNA following the manufacturer's instructions. The contraction rate of the

BAC was previously reported to be high [7], with rates between 20 % and up to 80 % depending on bacterial media and growth temperature conditions with richer media and faster/denser growth rates (Shaughn Bell, CSMC, personal communication). Therefore attention was paid to grow the cells in relative low density and at lower temperature (~ 27 °C).

#### Generation and culture of hiPSC lines from c9FTD/ALS patients

Human induced stem cells from a c9FTD/ALS patient were generated by transducing patient-derived fibroblasts with non integrating viral vectors (CytoTune 1.0 iPS Sendai Reprogramming Kit; Life Technologies) expressing the reprogramming factors Oct4, Sox2, Klf4 and c-Myc. Four weeks post transduction clones were manually picked and clonally expanded as hiPS cell lines on feeder cells. Established hiPSC cell colonies show the typical morphology of human pluripotent stem cells, stain positive for alkaline phosphatase (AP) and express the pluripotency-associated surface proteins TRA-1-60 and TRA-1-80. High resolution SNP-karyotyping was performed to exclude major karyotypic abnormalities induced by the reprogramming or culturing process (Suppl. Fig. 5 a-e).

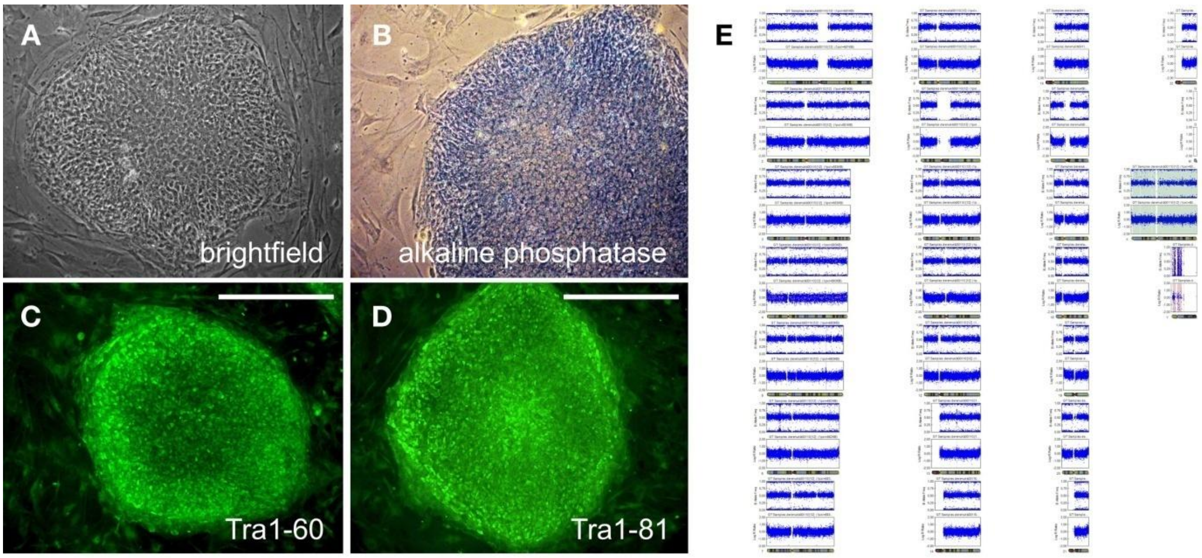


**Supplementary Figure 5 A - E: Characterization** of patient-derived hiPSC line 24/5#2

**A)** IPS cell colonies from cell line 24/5#2 show the typical morphology of human pluripotent stem cells, **B)** stain positive for alkaline phosphatase and express the pluripotency-associated surface proteins **C)** Tra1-60 and **D)** Tra1-80. **E)** High resolution SNP-karyotyping was performed to exclude major karyotypic abnormalities induced by the reprogramming process. Depicted are B-allele frequency and Log R ratio for each chromosome. Scale bars in C and D indicate 500 µm

For the study, the cultures were adapted to feeder free-culture conditions (mTeSR-E8, Stem Cell Technologies and BD-Matrigel, BD Biosciences) and passaged every 3-4 days as clumps with 0,5 mM EDTA in 1X PBS [16]. Detection of AP was performed using the VECTOR Blue Alkaline Phosphatase (Blue AP) Substrate Kit (Vector Laboratories) following the manufacturer's instructions. Staining against TRA-1-60 and TRA-1-81 was done as described below. Briefly cells were rinsed with 1X PBS followed by a paraformaldehyde (4 %) fixation for 10 min at RT. Samples were washed twice with 1X PBS prior incubation with primary antibodies against Tra1-60 and Tra1-81 (both 1:500 diluted, Thermo Fisher Scientific) for 2 h at RT. Cells were washed with 1X PBS, incubated with Alexa 488 anti-ms secondary antibody (1:1000, Thermo Fisher Scientific) and counterstained with DAPI (Sigma) for 45 min at RT in the dark. Samples were mounted in Mowiol 4-88 mounting solution (Carl Roth) for improved long term stability. High resolution SNP-karyotyping was performed as described previously [17].

Four hiPSC cell lines from three c9FTD/ALS patients were screened by Southern blot analysis (see below), and the line 24/5#2 (Suppl. Fig. 5 a-e) was selected for further nanopore sequencing analysis as it displayed at least two distinct maxima of the expanded allele around estimated 350 and 800 repeats.

Similarly, we adapted the previously characterized and described cell lines SC105iPS6 and SC105iPS7 from a FraX patient with concomitant Autism Spectrum Disorder to the same feeder-free culture and passaging conditions [18]. The cell lines were previously reported to carry approximately 380 (SC105iPS6) and 335 (SC105iPS7) CGG-repeats using the AmplideX FMR1 PCR Kit (Asuragen) according to the manufacturer's instruction [18]. Our Southern blot results showed peaks at approximately 750 (SC105iPS6) and 500 (SC105iPS7) CGG repeats (Suppl. Fig. 6) in agreement with the nanopore sequencing results from the same DNA preparations (see main text). We note that marked instability of FMR1-CGG STR expansions in other hiPSC and human embryonic stem cell lines through *in vitro* culture has been described before[19].

#### Clinical grade Southern blot-based determination of STR expansion size in DNA from C9FTD/ALS and FMR1 repeat expansion carrying hiPSC lines.

We used Southern blotting as means to estimate the repeat expansions present in our hiPSC lines from patients (Suppl. Fig. 6). The samples were processed and analyzed in the routine clinical diagnostic workflow for FXS and c9ALS/FTD (certified according to DIN EN ISO 15189:2014) at the Department for Human Genetics at Ulm University (Ulm, Germany). The diagnosticians were blinded for the experimental nature of the hiPSC samples and the DNA was processed in parallel with actual clinical samples.

Briefly, for the Southern blot determination of repeat expansion length of the $(G_4C_2)_n$- and $(CGG)_n$-repeats in our hiPSC lines 10 µg HMW DNA was digested overnight with HindIII (20 U) and XbaI [20 U, for $(G_4C_2)_n$] and EcoRI (20 U) and NruI [20 U, for $(CGG)_n$] respectively prior to electrophoresis. The fragmented gDNA was separated on a 0.8 % Agarose gel for 20 min with 180 V followed by 65 V overnight (C9orf72) or 20 min at 180 V followed by 60 V overnight (FMR1) respectively. The resulting gel was imaged with a ethidium bromide fluorescent stain and a copy of the image used for determination of a migration distance / DNA ladder standard curve.

DNA was transferred to a positively charged nylon membrane (Roche Applied Science) by capillary blotting and was baked at 80 °C for 2 hr.

The hybridization probes were either a 210 bp PCR-fragment located upstream of the $G_4C_2$-repeat in the *C9orf72*-gene or a 480 bp PCR-fragment located downstream of the CGG-Repeat in the *FMRI*-gene respectively. Probes (100 ng/ filter) were labelled with 50 mCi $\alpha^{32}$PdCTP and hybridized to the filters at 71 °C overnight. After washing, X-ray films were exposed to the filters. The lengths of hybridizing fragments were calculated in relation to λ-DNA digested with BstEII. Fragments derived from wildtype *C9orf72*-alleles are approximately 2.3 kb in length. Wildtype *FMRI*-fragments from active, unmethylated X-chromosomes are approximately 2.9 kb, from inactivated, methylated X-chromosomes 5.2 kb in length as the NruI restriction enzyme used in this assay cuts only 5'-TCGCGA-3' sequences in which the CpGs are unmethylated.

**Supplementary Table 1: Cas12a** crRNAs and bottom strand (BS) barcodes used in this study.

| ROI | crRNA ID | crRNA name | ref (hg38) | strand | protospacer 5'->3' | overhang | BS barcode 5'->3' |
|---|---|---|---|---|---|---|---|
| FMR1 | FMR1-C1 | FMR1_Cpf1_147913435_C1 | chrX: 147913437-147913457 | - | CCAGCCTTCCTTCCACACGCA | AGGT_- | /5Phos/AGGTGGTGCTGAAGAAAGTTGTCGGTGTCTTTGTGTTAACCTTAGCAAT |
| FMR1 | FMR1-C2 | FMR1_Cpf1_147916117_C2 | chrX: 147916120-147916140 | - | TAACTTTATCTTTCCTTAACA | CCTG_- | /5Phos/CCTGGGTGCTGAAGAAAGTTGTCGGTGTCTTTGTGTTAACCTTAGCAAT |
| FMR1 | FMR1-C3 | FMR1_Cpf1_147908315_C3 | chrX: 147908319-147908339 | + | ATGGAAACCAAGGGCCAAGGC | CTGC_+ | /5Phos/CTGCGGTGCTGAAGAAAGTTGTCGGTGTCTTTGTGTTAACCTTAGCAAT |
| FMR1 | FMR1-C4 | FMR1_Cpf1_147910463_C4 | chrX: 147910467-147910487 | + | AGCCCTATTGGGTTCTTGGCC | GAGG_+ | /5Phos/CTGCGGTGCTGAAGAAAGTTGTCGGTGTCTTTGTGTTAACCTTAGCAAT |
| FMR1 | FMR1-C5 | FMR1_Cpf1_147911947_C5 | chrX: 147911951-147911971 | + | ACTTCCGGTGGAGGGCCGCCT | GAGA_+ | /5Phos/GAGAGGTGCTGAAGAAAGTTGTCGGTGTCTTTGTGTTAACCTTAGCAAT |
| C9orf72 | C9orf72-C1 | C9orf72_Cpf1_27571873_C1 | chr9: 27571877-27571897 | + | CAGTACCAGAAAGTTCACAAC | GTGT_+ | /5Phos/GTGTGGTGCTGAAGAAAGTTGTCGGTGTCTTTGTGTTAACCTTAGCAAT |
| C9orf72 | C9orf72-C2 | C9orf72_Cpf1_27571960_C2 | chr9: 27571964-27571984 | + | TCACAGTTCCAAGTTTCTCAG | GTCT_+ | /5Phos/GTCTGGTGCTGAAGAAAGTTGTCGGTGTCTTTGTGTTAACCTTAGCAAT |
| C9orf72 | C9orf72-C3 | C9orf72_Cpf1_27574577_C3 | chr9: 27574580-27574600 | - | TTCCTCCCTTTCTTCCTCGGT | TCAC_- | /5Phos/TCACGGTGCTGAAGAAAGTTGTCGGTGTCTTTGTGTTAACCTTAGCAAT |
| C9orf72 | C9orf72-C4 | C9orf72_Cpf1_27573666_C4 | chr9: 27573669-27573689 | - | CCACCCTCTCTCCCCACTACT | CAAG_- | /5Phos/CAAGGGTGCTGAAGAAAGTTGTCGGTGTCTTTGTGTTAACCTTAGCAAT |
| C9orf72 | C9orf72-C5 | C9orf72_Cpf1_27572279_C5 | chr9: 27572283-27572303 | + | CTAAAGTGGCAGGCCTTGGCA | TCTG_+ | /5Phos/TCTGGGTGCTGAAGAAAGTTGTCGGTGTCTTTGTGTTAACCTTAGCAAT |
| C9orf72 | C9orf72-C6 | C9orf72_Cpf1_27572805_C6 | chr9: 27572809-27572829 | + | AGCAAGTCTGTGTCATCTCGG | CTCC_+ | /5Phos/CTCCGGTGCTGAAGAAAGTTGTCGGTGTCTTTGTGTTAACCTTAGCAAT |

**Supplementary Table 2:** Overview of oxford nanopore sequencing runs performed in this study.

| | | | | | i | | ii | iii | iv | v | vi | vii |
|---|---|---|---|---|---|---|---|---|---|---|---|---|
| **ID** | **Kit** | **input sample** | **ROI** | **crRNA ID** | **ROI** | | **D** | **F** | **S** | **EF** | **E** | **W** |
| | | | | | crRNA | ROI size | reads on target | Fraction of ROI covered | ROI reads/ all mapped reads (x10e-6) | D for ROI / D genome | exp/wt | input DNA ug |
| FAH31667 | SQK-LSK108 | plasmid | NA | | | NA | 2491356 | | | | | |
| FAE22321 | SQK-RAD001 | BAC | NA | | | NA | 89103 | | | | | |
| FAH31320 | SQK-LSK108 | C9orf72 | C9ORF72 | none | | NA | 1 | NA | 0,055 | 0,271 | 0,000 | 2 |
| FAH37180 | SQK-LSK108 | C9orf72 | C9ORF72 | none | | NA | 5 | NA | 0,442 | 2,136 | 0,000 | 2 |
| FAH34894 | SQK-LSK108 | C9orf72 | C9ORF72 | none | | NA | 7 | NA | 0,361 | 1,782 | 0,000 | 2 |
| FAH33354 | SQK-LSK108 | C9orf72 | C9ORF72 | none | | NA | 0 | NA | 0,000 | 0,000 | | 2 |
| FAH42748 | SQK-LSK108 | C9ORF72 24/5#2 | C9ORF72 | C9orf72-C4, C5 | c4-c5 | 1369 | 1 | NA | 56,818 | 145,954 | Inf | 10 |
| FAH35601 | SQK-LSK108 | C9ORF72 24/5#2 | C9ORF72 | C9orf72-C4, C5 | c4-c5 | 1369 | 5 | NA | 121,465 | 212,306 | 0,333 | 10 |
| FAH67301 | SQK-LSK108 | C9ORF72 24/5#2 | C9ORF72 | C9orf72-C2, C4 | c2-c4 | 1688 | 10 | NA | 909,008 | 4678,266 | 0,143 | 10 |
| FAH65171 | SQK-LSK108 | FMR1 SC105iPS7 | FMR1 | FMR1-C3 | | NA | 13 | NA | 71,122 | 352,875 | | 10 |
| FAH66274 | SQK-LSK108 | FMR1 SC105iPS6 | FMR1 | FMR1-C1, C3 | c1-c3 | 5102 | 2 | NA | 131,570 | 274,935 | | 10 |
| FAH81424 | SQK-LSK108 | C9ORF72 24/5#2 | C9ORF72 | C9orf72-C3, C4 | | NA | 1 | NA | 70,343 | 162,703 | 0,000 | 10 |
| FAH72446 | SQK-LSK108 | C9ORF72 24/5#2 | C9ORF72 | C9orf72-C3, C4 | | NA | 9 | NA | 269,211 | 757,130 | 0,400 | 10 |
| FAH67421 | SQK-LSK108 | C9ORF72 24/5#2 | C9ORF72 | C9orf72-C4 | | NA | 21 | NA | 148,567 | 226,106 | 0,273 | 10 |
| FAH80338 | SQK-LSK108 | C9orf72 24/5#2 | C9ORF72 | C9orf72-C1,C4 | c1-c4 | 1775 | 2 | NA | 45,087 | 63,997 | 0,000 | 10 |
| FAH81473 | SQK-LSK108 | C9orf72 24/5#2 | C9ORF72 | C9orf72-C1,C4 | c1-c4 | 1775 | 3 | NA | 55,989 | 73,985 | Inf | 10 |
| FAH87258 | SQK-LSK109 | C9orf72 24/5#2 | C9ORF72 | C9orf72-C4 | | NA | 8 | NA | 10,864 | 36,514 | 0,600 | 5 |
| FAH88643 | SQK-LSK109 | C9orf72 24/5#2 | C9ORF72 | C9orf72-C4 | | NA | 11 | NA | 37,748 | 111,099 | 0,333 | 5 |
| FAK02017 | SQK-LSK109 | SC105iPS7 | FMR1 | FMR-C1, C3 | c1-c3 | 5102 | 27 | NA | 5,034 | 27,238 | | 5 |
| FAJ02524 | SQK-LSK109 | C9ORF72 24/5#2 | C9ORF72 | C9orf72-C1, C2, C3, C4 | c2-c4 | 1688 | 118 | NA | 22,844 | 125,931 | 0,466 | 5 |
| FAH91937 | SQK-LSK109 | C9ORF72 24/5#2 | C9ORF72 | C9orf72-C1, C2, C3, C4 | c2-c4 | 1688 | 72 | NA | 18,393 | 96,494 | 0,784 | 5 |
| FAJ02378 | SQK-LSK109 | SC105iPS6 | C9ORF72 | C9orf72-C1, C2, C3, C4 | c2-c4 | 1688 | 350 | NA | 67,460 | 363,316 | 0,000 | 5 |
| | | | FMR1 | FMR1-C3 | | NA | 4 | NA | 0,771 | 4,152 | | |
| FAH66294 | SQK-LSK109 | SC105iPS6 | C9ORF72 | C9orf72-C1, C2, C3, C4 | c2-c4 | 1688 | 28 | NA | 23,024 | 121,619 | 0,048 | 5 |
| | | | FMR1 | FMR1-C3 | | NA | 1 | NA | 0,822 | 4,344 | | |
| PAD01034 | SQK-LSK109 | SC105iPS6 | C9ORF72 | C9orf72-C1, C2, C3 | c2-c3 | 259 | 164 | NA | 108,023 | 439,041 | 0,009 | 5 |
| | | | FMR1 | FMR1-C3 | | NA | 3 | NA | 1,976 | 8,031 | | |
| PAD01039 | SQK-LSK109 | C9ORF72 24/5#2 | C9ORF72 | C9orf72-C1, C2, C3 | c2-c3 | 2599 | 347 | NA | 44,980 | 196,485 | 0,455 | 5 |
| | | | FMR1 | FMR1-C3 | | NA | 25 | NA | 3,241 | 14,156 | | |
| PAD01413 | SQK-LSK109 | SC105iPS6 | C9ORF72 | C9orf72-C1, C2, C3 | c2-c3 | 2599 | 804 | NA | 96,546 | 427,447 | 0,002 | 5 |
| | | | FMR1 | FMR1-C1, C2, C3, C4 | c1-c4 | 2954 | 316 | NA | 37,946 | 168,001 | | |
| PAD01249 | SQK-LSK109 | C9ORF72 24/5#2 | C9ORF72 | C9orf72-C1, C2, C3 | c2-c3 | 2599 | 600 | NA | 76,558 | 327,570 | 0,403 | 5 |
| | | | FMR1 | FMR1-C1, C2, C3, C4 | c1-c4 | 2954 | 691 | NA | 88,170 | 377,251 | | |

| | | |
|---|---|---|
| **(i)** | **Region of interest (size): $ROI_{wt}$;** | here we report the fragment size of the wild type allele between the two most proximal crRNA binding sites |
| **(ii)** | **Total average read depth (in ROI): D;** | here we report the number of reads spanning the repeat expansion site |
| **(iii)** | **Fraction of ROI sufficiently covered (at a specified D): F;** | NA |
| **(iv)** | **Specificity (fraction of reads in ROI): S;** | here we report all reads in ROI versus all reads (mappable reads which is excluding adapters) obtained by a single flow cell |
| **(v)** | **Enrichment Factor (D for ROI versus D): EF;** | D for ROI versus D for rest of genome |
| **(vi)** | **Evenness (lack of bias): E;** | here we report in case of two alleles with one wild type and the other expanded the ratio expanded allele over the wild type allele |
| **(vii)** | **Weight (input DNA requirement): W** | here we report the actual amount of DNA used for a specific experiment |

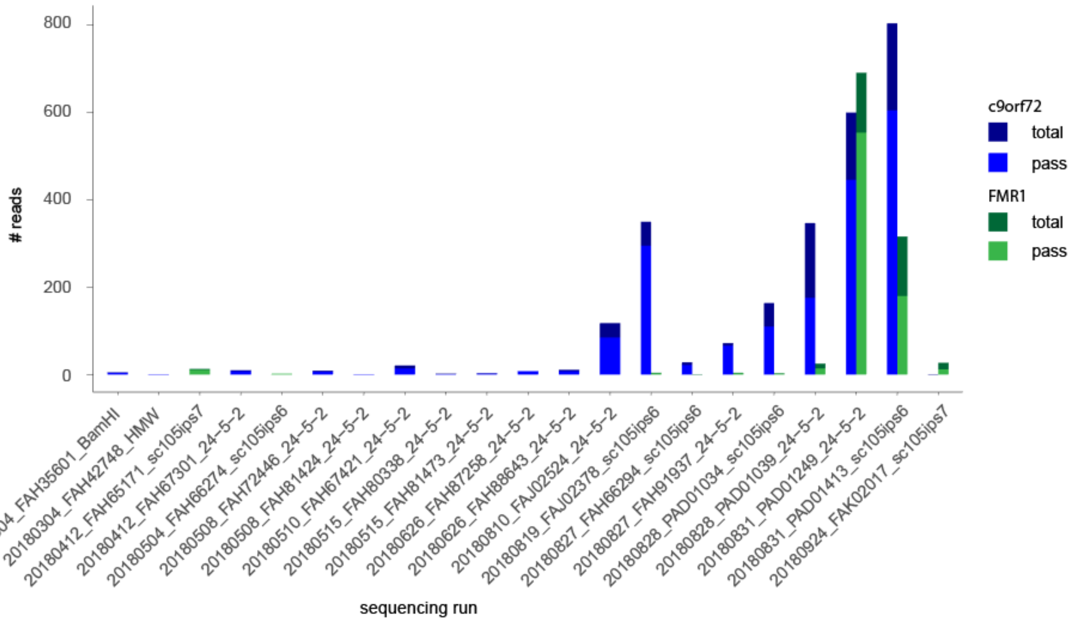


**Supplementary Figure 6:** Sequencing throughput

Number of reads on target and number of reads passing the STRique pipeline for all Flow-Cells used in this study.

**Supplementary Table 3:** Oligomere sequences used for synthetic $(G_4C_2)_n$-repeat cloning.

| Oligo ID | Oligo sequence 5'→3' |
|---|---|
| 3x GGGGCC fwd | GGATCCGCTCTTCCGGGGCCGGGGCCGGGGCCGGGGCCTGCGGCCGC |
| 3x GGGGCC rev | GCGGCCGCAGGCCCCGGCCCCGGCCCCGGCCCCGGAAGAGCGGATCC |
| 4x GGGGCC fwd | GGATCCGCTCTTCCGGGGCCGGGGCCGGGGCCGGGGCCGGGGCCTGCGGCCGC |
| 4x GGGGCC rev | GCGGCCGCAGGCCCCGGCCCCGGCCCCGGCCCCGGCCCCGGAAGAGCGGATCC |

#### Code availability

All custom code developed for this study is under MIT license available at: https://github.com/giesselmann/STRique

The RepeatHMM package was forked and is modified available at:

https://github.com/giesselmann/RepeatHMM

This study used previously published software: basecalling was performed with ONT Albacore 2.3.3, alignment was performed with minimap2 (v2.12), methylation analysis was performed with graphmap (v0.5.2) and nanopolish (v0.9.0). for raw nanopore signal processing seqan (v2.4.0) was used.

#### Data availability

plasmid 10k random sample per barcode (fast5 + fasta)

BAC reads mapping to repeat (fast5 + fasta)

human wgs: Basecalled reads (fasta) reads on target (fast5)

human enrichment: mapped reads (fasta) reads on target (fast5)

1. Reinert, K. *et al.* The SeqAn C++ template library for efficient sequence analysis: A resource for programmers. *J. Biotechnol.***261,** 157–168 (2017).
2. Schreiber, J. & Karplus, K. Analysis of nanopore data using hidden Markov models. *Bioinformatics***31,** 1897–1903 (2015).
3. Jain, M. *et al.* Nanopore sequencing and assembly of a human genome with ultra-long reads. *Nature Biotechnology***36,** 338–345 (2018).
4. Liu, Q., Zhang, P., Wang, D., Gu, W. & Wang, K. Interrogating the 'unsequenceable' genomic trinucleotide repeat disorders by long-read sequencing. *Genome Medicine 2017 9:1***9,** 65 (2017).
5. Dashnow, H. *et al.* STRetch: detecting and discovering pathogenic short tandem repeat expansions. *Genome Biol***19,** 121 (2018).
6. Sareen, D. *et al.* Targeting RNA Foci in iPSC-Derived Motor Neurons from ALS Patients with a C9ORF72 Repeat Expansion. *Sci Transl Med***5,** 208ra149 (2013).
7. O'Rourke, J. G. *et al.* C9orf72 BAC Transgenic Mice Display Typical Pathologic Features of ALS/FTD. *Neuron***88,** 892–901 (2015).
8. Sambrook, J. & Maniatis, T. *Molecular Cloning*. (Cold Spring Harbor Laboratory Press, 1989).
9. Montague, T. G., Cruz, J. M., Gagnon, J. A., Church, G. M. & Valen, E. CHOPCHOP: a CRISPR/Cas9 and TALEN web tool for genome editing. *Nucl. Acids Res.***42,** W401–W407 (2014).
10. Labun, K., Montague, T. G., Gagnon, J. A., Thyme, S. B. & Valen, E. CHOPCHOP v2: a web tool for the next generation of CRISPR genome engineering. *Nucl. Acids Res.***44,** W272–W276 (2016).
11. Zetsche, B. *et al.* Cpf1 Is a Single RNA-Guided Endonuclease of a Class 2 CRISPR-Cas System. *Cell***163,** 759–771 (2015).
12. Strohkendl, I., Saifuddin, F. A., Rybarski, J. R., Finkelstein, I. J. & Russell, R. Kinetic Basis for DNA Target Specificity of CRISPR-Cas12a. *Mol Cell***71,** 816–824.e3 (2018).
13. Chen, J. S. *et al.* CRISPR-Cas12a target binding unleashes indiscriminate single-stranded DNase activity. *Science***360,** 436–439 (2018).
14. Mizielinska, S. *et al.* C9orf72 repeat expansions cause neurodegeneration in Drosophila through arginine-rich proteins. *Science***345,** 1192–1194 (2014).
15. Kwok, Y. K. *et al.* Validation of a robust PCR-based assay for quantifying fragile X CGG repeats. *Clinica Chimica Acta***456,** 137–143 (2016).
16. Chen, G. *et al.* Chemically defined conditions for human iPSC derivation and culture. *Nat Methods* (2011). doi:10.1038/nmeth.1593
17. Mertens, J. *et al.* APP processing in human pluripotent stem cell-derived neurons is resistant to NSAID-based γ-secretase modulation. *Stem Cell Reports***1,** 491–498 (2013).
18. Boland, M. J. *et al.* Molecular analyses of neurogenic defects in a human pluripotent stem cell model of fragile X syndrome. *Brain***140,** 582–598 (2017).
19. Zhou, Y., Kumari, D., Sciascia, N. & Usdin, K. CGG-repeat dynamics and FMR1 gene silencing in fragile X syndrome stem cells and stem cell-derived neurons. *Molecular Autism***7,** 165 (2016).
