## Supplementary material for "Repeat expansion and methylation state analysis with nanopore sequencing"

### Supplementary Figure 6

#### Figure Legend

**Supplementary Figure 6a**
Autoradiographic Southern blot from controls and C9orf72 expanded allele carriers. Uncropped, unmodified scan of a Southern blot analysis of the cell line used for this study.

**Supplementary Figure 6b**
Autoradiographic Southern blot from controls and C9orf72 expanded allele carriers with labels. Same scan of a Southern blot analysis of the cell line used for this study as in Suppl. Fig 6a with labels indicating the sample names and function (e.g. negative control, mutation carrier). Legend: hiPSC = human induced pluripotent stem cell; HMW-DNA = high molecular weight DNA; -m = methylated allele; wt = wildtype; yellow text/lines = hiPSC lines used in this study

**Supplementary Figure 6c**
Autoradiographic Southern blot from controls and FMR1 expanded allele carriers. Uncropped, unmodified scan of a Southern blot analysis of the cell line used for this study.

**Supplementary Figure 6d**
Autoradiographic Southern blot from controls and FMR1 expanded allele carriers with labels. Same scan of a Southern blot analysis of the cell line used for this study as in Suppl. Fig 6e with labels indicating the sample names and function (e.g. negative control, mutation carrier). Fragments covering the FMR1 CGG-STR migrate in this assay depending on their methylation status: methylated fragments as the NruI restriction enzyme does not cut if its 5'-TCGCGA-3' restriction site is methylated at the CpG site. As a result, the restriction fragment is longer and spans from the two adjacent HindIII restriction sites instead from the HindIII – NruI sites. Legend: hiPSC = human induced pluripotent stem cell; HMW-DNA = high molecular weight DNA; -m = methylated allele; wt = wildtype; -u = unmethylated allele; yellow text/lines = hiPSC lines used in this study

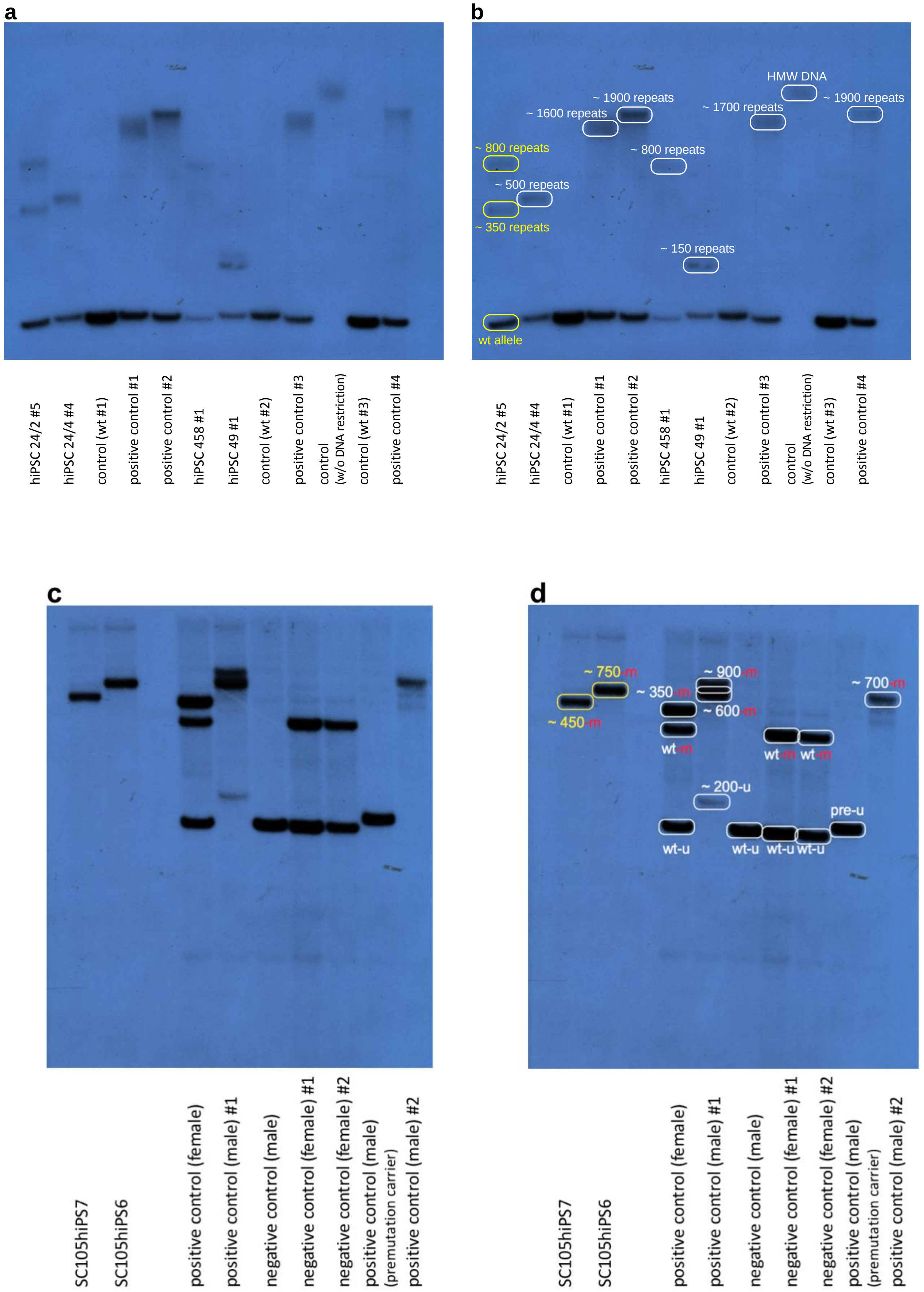


**Supplementary Figure 6**
