## Supplementary material for "Repeat expansion and methylation state analysis with nanopore sequencing"

* corresponding author

### 1 Considerations pertaining to the nomenclature, selection of prototypical examples for unstable repeat expansions.

In our study, we set out to explore this technology as means to specifically sequence selected genomic regions, to ascertain repeat expansion status and to determine CpG methylation patterns on the single DNA molecule level from cells from c9FTD/ALS and FXS patients as prototypical examples for STR expansion disorders.

We acknowledge that several, at times contradictory, abbreviations and names are used in the literature for unstable repeat expansions in health and disease. In this paper, we chose to follow the nomenclature of the Medical Subject Header (MeSH) descriptor data 2018 structure for Tandem Repeat Sequences [G05.360.340.024.850, https://meshb.nlm.nih.gov/record/ui?ui=D020080]: We use the term 'Short Tandem Repeats' (STRs) interchangeably with "Microsatellite Repeats" to refer to short repeat units of 2-8 base pairs that are tandemly repeated (see Suppl. Fig. 7). Longer repeats of 10 or more repeats are referred to as "Variable Nucleotide Tandem Repeats" (VNTR) (see Suppl. Figure 7), for which MeSH suggested the alternative term „minisatellite repeats"[G05.360.340.024.850.550]. Visual representations of n-tuple sequence motifs are presented in Suppl. Fig. 7

Notably, several recent studies defined STRs as consisting of 1-6 bp motifs (see for example Hannan, Nat Rev Genetics 2018[1] and Dashnow et al. Genome Biology 2018[2]); thus not in line with the definition by MeSH and we do acknowledge that not all MeSH descriptors may reflect current consensus in the field or expert knowledge and may lack comprehensiveness (e.g. tandem repeats with nine base pairs as repetitive motiv - following the MeSH descriptors - are neither microsatellites nor minisatellites nor STRs nor VNTRs). Yet we chose the MeSH ontology as actively curated, maintained and publicly available resource for a systematic view on repetitive DNA sequences.

We wish to point out that, STRs do not only occur in disease state (for recent examples see [2-4]) but also occur in the physiological statein the context of so-called microsatellites, numerous polymorphic genomic loci which are not necessarily associated with any disorder or pathology [5].

Both methods described in our study are generally applicable to any genomic region (CRISPR Cas12a-RNP enrichment) or any STR (nanoSTRIque) independently of their potential pathogenic role.

**Supplementary Figure 7: Systematic overview over tandem repeats and their nomenclature**

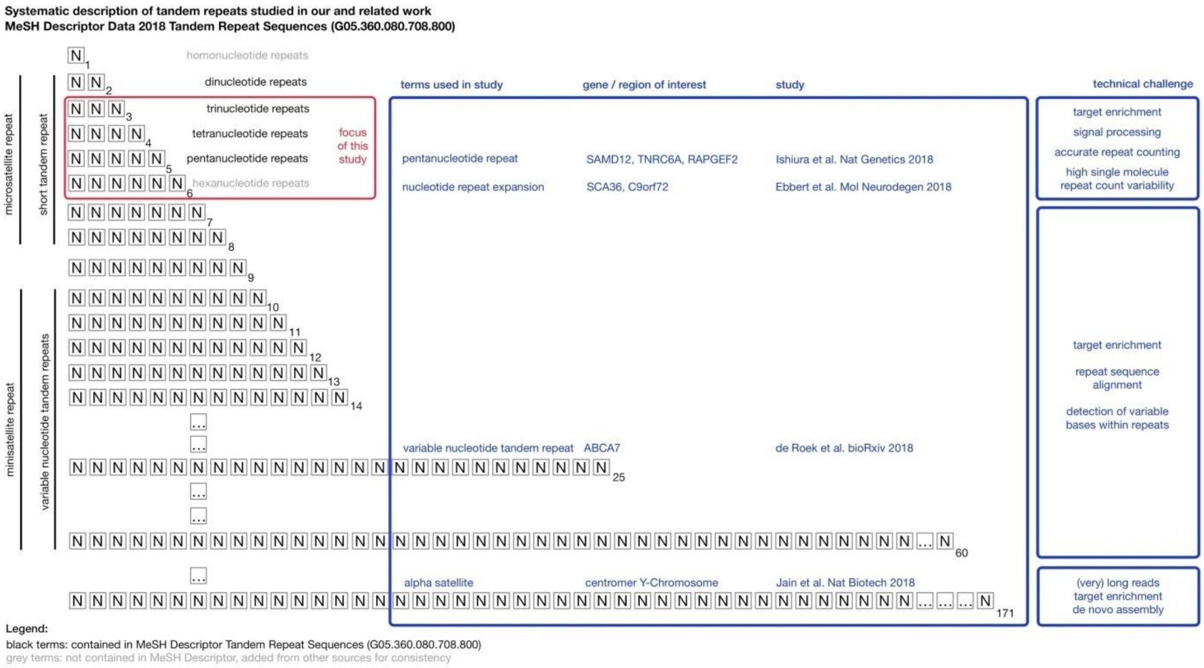


For context, we have added all four currently available studies (2018/10/15) that have used nanopore sequencing to characterize different tandem repeats on the background of the human genome and added the terms used in these studies in the context of MeSH descriptors.On the right side of Suppl. Figure 7 we list technical challenges specific to the application of nanopore sequencing to the different subsets of tandem repeats.
Importantly, different repeat expansion classes pose different technical problems from the perspective of using third generation sequencing technologies (Suppl. Table 4). In our study we experimentally focussed on STRs with 3-6 bp repetitive elements as these pose a nanopore sequencing and signal processing challenge distinct from those associated with VNTRs/minisatellites or even larger tandem repeat structures such as centromeric repeats and will accommodate for most currently known and clinically relevant tandem repeat expansion associated disorders.
Briefly, nanopore signals from 3-6 bp STRs deviate strongly from many assumptions and parameter sets used for general purpose base callers (such as ONT's Albacore). For example, asdemonstrated in Fig. 1A and Suppl. Fig. 2, raw signals from repeats are much more homogenous and often are of smaller amplitude when compared to more variable sequences. In comparison, the much larger VNTRs/minisatellites (10 or more bp) can be effectively basecalled with general purpose base callers such as Albacore at the disadvantage of high base-calling error rates in single reads of 10-15 %. This poses a problem for the accurate determination of repeat expansion counts[6] different from those encountered with shorter repetitive motifs. Very large tandem repeat units, on the other hand, can be more effectively base-called and sequence variants can be detected through raw signal polishing methods but reads spanning several hundreds of kilobases are necessary to correctly assemble such higher order repetitive sequences.
From a clinical perspective, unstable genomic short repetitive sequences (STRs with 3-6 bp) cause more than 30 Mendelian human disorders[1,7,8]. In this study, we employ FXS and c9FTD/ALS al prototypical examples for clinically relevant repeat expansion disorders in

humans to demonstrate the general applicability and potential usefulness of the combination of both novel methods towards an integrative analysis of unstable repeat expansions for basic research and clinical applications.

We chose the GGGGCC hexanucleotide [$(G_4C_2)$n] repeat expansion in the C9orf72 gene as a example for a hexanucleotide repeat expansion disorders, as it represents the first and most common known genetic variant underlying both Amyotrophic Lateral Sclerosis and Frontotemporal Dementia (c9FTD/ALS)[9,10]. Based on Southern blot analyses, (G4C2)n-expansions in c9FTD/ALS patients have been estimated in the range of 100-7500 repeats resulting in 0.6 to 45 kB regions with 100 % Guanine/Cytosine content and G-quadruplex formation[11-13]. These sequence features and structural DNA variants are often mentioned as reason, why conventional, polymerase-based molecular methods fail to resolve repeat expansion numbers higher than 100 - 150 repeats[9,10,13,14].

We also chose Fragile X Syndrome (FXS) as second example for a STR expansion disorder. In FXS, as it represents the first described repeat expansion disorder, the accumulation of more than 200 CGG-repeats impair the function of the FMR1-gene and subsequently result in neurodevelopmental delay, intellectual disability and frequently autism[15].

In both disorders, STR expansions are intra- and interindividually highly variable due to genetic anticipation, ageing, neural differentiation and epigenetic mechanisms. Of these, genetic anticipation has been well described for both c9FTD/ALS and FXS.[1,16] Genetic anticipation describes the observation that disease severity in most if not all STR expansion disorders increases through successive generations, i.e. with progressively earlier clinical onset and more severe phenotypes. Genetic anticipation is now mainly ascribed to the fact that expanded repeats dynamically increase in length over generations and that longer repeats tend to be increasingly unstable; and longer repeats are associated with more severe phenotypes. In the case of c9FTD/ALS longer STR lengths have been associated with earlier age of disease onset,higherdisease severity and variation between tissues and affected brain regions[12,17-22].

In both c9FTD/ALS and FXS, increased methylation of CpG islands close to the expanded STRs have been also implicated to affect disease phenotype[19,20,23-25].

We note, that due to the focus of this study on the universal nature of our proposed approach and inherent space constraints, we can only superficially describe the clinical presentation and specific aspects of both disorders, such as premuation status, associated disorders such as Fragile X associated tremor/ataxia syndrome (FXTAS).

In this discussion, we therefore highlight only aspects of both disorders pertinent to the interpretation and novel insights enabled by the methods. We encourage interested readers to consult existing, excellent reviews[1,8,16] as a starting point for a much more comprehensive summary of unstable repeat expansion disorders in general and c9FTD/ALS and FXS in particular.

### 2 Considerations pertaining to the current challenges for exact repeat expansion quantification with polymerase based methods, Southern blotting and third generation sequencing methods

As of now, most methods for the determination of repeat expansion status each suffer from at least two of the following three problems: i) lack of resolution at higher repeat numbers, i) lack of single molecule analysis capability and/or iii) targeting of expansion loci for sufficient coverage of higher repeat expansion counts.
Because of the limitations of each currently available methods, the combination of PCR and Southern blots is still recommended in the diagnostic setting (e.g. for FXS).[26,27]

Conventional, polymerase-based methods (PCR-reactions, Sanger-sequencing) are currently frequently used to estimate repeat numbers, but are imperfect beyond the threshold of 100-150 repeats[2,10,28] as DNA polymerases e.g. in conventional PCR or sequencing reactions cannot read through the repetitive, GC-rich and often secondary structures forming sequences. Although, while few heterogeneous alleles with lower repeat counts can be detected with PCR-methods, this readout degrades quickly with higher repeat counts > 100. Also, no conventional polymerase-based methods for repeat expansion status on the single molecule level is currently available.

Short read sequencing technologies (mainly Illumina®) are limited by their read/insert length[29] as even paired end reads can span only smaller repeat expansions.Synthetic long read methods coupled with Illumina short read sequencing also fail to resolve longer repeat expansions because the mapping problem with sequences containing 100 % repetitive elements remain.

Southern blot analysis requires large amounts of DNA and cannot be scaled down to the single molecule level. Additionally, the resolution of gel electrophoresis analysis decreases strongly with repeat expansion length (see for example Suppl. Fig. 5a - h and here specifically the standard curves Suppl. Fig. 5d for the c9FTD/ALS and Suppl. Fig. 5h). We note, that only Southern blots are currently realistically capable of resolving intraindividual repeat instability/heterogeneity if different distinct high repeat number expansion maxima exist as has been discussed e.g. in Almeida et al. 2013.[30]

Current third generation single molecule sequencing methods (PacBio SMART-seq and ONT nanopore sequencing) have been consistently shown to be capable to read through long stretches of repetitive DNA from tandem repeat expansion disorder derived plasmids[31-34], but their throughput from patient samples has been up to now severely limited.
As illustrative example, Ebbert and colleagues[32] recently reported the results from a whole genome SMRT-seq experiment employing five PacBio flow cells with a total of four PacBio reads covering C9orf72 repeat expansion region (30-1324 repeats). Only one read with 1324 repeat counts fully bridged the repeat expansion region. A targeted enrichment experiment utilizing the “No-Amp” Cas9-targeting SMRT-seq technology[35] in the same study resulted in a total of 134 reads from expanded alleles with a repeat expansion count distribution (two modes at 110 and 870), which were interestingly not matching results obtained from Southern blot analysis from the same subject (around estimated 1000 repeats in the Southern blot). Ebbert et al. also reported the results from nanopore whole genome sequencing experiments with 2 reads from the wild type allele and no single read covering

the C9orf72 repeat expansion could be recovered from 15 flow cells loaded with whole genome libraries.

Supplementary Table 4: Comparison of methods for the determination of repeat expansion status and quantification of repeat expansion counts

| **Technology** | **targeting strategy** | **resolution** | **singlemolecule** | **targeting/ coverage** |
|---|---|---|---|---|
| Southern blot | hybridization probe | + low rn<br>(+) high rn | no | +++ |
| repeatprimed PCR | PCR primer | ++ lowrn<br>- high rn | no | +++ |
| Sanger | Sequencing primer | ++ low rn<br>- high rn | no | +++ |
| Illumina short read | n.a. | +++ low rn<br>- high rn | no | +++ |
| PacBioNo-Amp SMRT-seq | Cas9-targeting of region, hairpin adaptor ligation and biotin pulldown | +++ low rn<br>(+) high rn | yes | (+) |
| Nanopore Cas12a-RNP* | Cas12a-RNP targeting, and selective adaptor ligation | +++ low rn<br>+++ high rn | yes | +++ |

Legend: * this study; rn, repeat number

### 3 Considerations pertaining to reporting enrichment results

In this study we report our enrichment results in reads per target per individual flow cell which equals a single DNA sample from a single individual. From this metric, we derive sub-counts, e.g. reads covering the target with or without repeat expansion and from those, which appear to contain a repeat expansion, those, from which we could derive a repeat count with nanoSTRIque.

We believe that this metric captures best the real world results, where a diagnostician or researcher analyzes a sample from an individual to learn about the length of a possible repeat expansion and its epigenetic context in one flow cell. For consistency reasons we chose to report only these results in the main text.

We note that a set of several other metrics have been proposed previously to describe targeted sequencing enrichment results[36]. The parameters proposed were adapted to our specific experimental setup and research question as described below the parameter description written in bold.

- **(i) Region of interest (size): $ROI_{wt}$;**
  here we report the fragment size of the wild type allele between the two most proximal crRNA binding sites
- **(ii) Total average read depth (in ROI): D;**
  here we report the number of reads spanning the repeat expansion site
- **(iii) Fraction of ROI sufficiently covered (at a specified D): F;**
  NA
- **(iv) Specificity (fraction of reads in ROI): S;**
  here we report all reads in ROI versus all reads (mappable reads which is excluding adapters) obtained by a single flow cell
- **(v) Enrichment Factor (D for ROI versus D): EF;**
  D for ROI versus D for rest of genome
- **(vi) Evenness (lack of bias): E;**
  here we report in case of two alleles with one wild type and the other expanded the ratio expanded allele over the wild type allele
- **(vii) Weight (input DNA requirement): W**
  here we report the actual amount of DNA used for a specific experiment

The following table exemplarily summarizes these metrics applied to our two target regions and the maximum enrichment results obtained with the methods from single flow cells.
Results from all flow cells following the above metrics are listed in Supplementary Table 2.

### 4 Considerations pertaining to the determination of CpG methylation on unstable repeat expansions in nanopore reads

In this study we provide evidence for the ability of our proposed workflow to detect CpG methylation on the single molecule level with nanopolish[37] and to connect this information with repeat expansion counts from nanoSTRIque. It is important to note, that nanopolish fails to call CpG methylation of the repeat expansion itself (data not shown). This challenge will be likely difficult to overcome in the near future, as the raw nanopore current differences induced by cytosine methylation are on the range of 1/10 of the amplitude changes resulting from any other base passing the pore[37-39]. The mCpG signal can be readily detected in about 75 % of all CpGs on the single read level, but successful CpG methylation determination strongly depends on the bioinformatic normalization of the raw current signals to correct for several noise parameters independent of the base composition and cytosine-modification status.

We found two major reasons for not being able to obtain methylation readout from the repeat sequence itself. The first one is the base-calling error within the repeat making the sequence alignment input of nanopolish even with a repeat count corrected reference less reliable.The second reason is the internal grouping of adjacent CpGs within nanopolish providing group methylation prediction for CpG sites in close proximity to each other. In the case of the $(G_4C_2)_n$ repeat we assume the formation of such a single methylation prediction group exceeding memory and runtime limits.

Therefore we restricted our DNA modification analysis to the regions surrounding the repeats and controlled for possible effects of the repeat expansion on the base calling and cytosine modification detection with plasmid/BAC DNA from bacterial cells with known methylation patterns (Suppl. Fig. 4).

**References:**


1. Hannan, A. J. Tandem repeats mediating genetic plasticity in health and disease. *Nat Rev Genet***19,** 286–298 (2018).
2. Dashnow, H. *et al.*STRetch: detecting and discovering pathogenic short tandem repeat expansions. *Genome Biol***19,** 121 (2018).
3. Sun, J. H. *et al.* Disease-Associated Short Tandem Repeats Co-localize with Chromatin Domain Boundaries. *Cell***175,** 224–238.e15 (2018).
4. Gymrek, M. A genomic view of short tandem repeats. *Curr. Opin. Genet. Dev.***44,** 9–16 (2017).
5. Dib, C. *et al.* A comprehensive genetic map of the human genome based on 5,264 microsatellites. *Nature***380,** 152–154 (1996).
6. De Roeck, A. *et al.* Accurate characterization of expanded tandem repeat length and sequence through whole genome long-read sequencing on PromethION. *bioRxiv* 439026 (2018). doi:10.1101/439026
7. Essebier, A. *et al.* Statistical Enrichment of Epigenetic States Around Triplet Repeats that Can Undergo Expansions. *Front. Neurosci.***10,** 92 (2016).
8. Gatchel, J. R. &Zoghbi, H. Y. Diseases of unstable repeat expansion: mechanisms and common principles. *Nat Rev Genet***6,** 743–755 (2005).
9. DeJesus-Hernandez, M. *et al.* Expanded GGGGCC hexanucleotide repeat in noncoding region of C9ORF72 causes chromosome 9p-linked FTD and ALS. *Neuron***72,** 245–256 (2011).
10. Renton, A. E. *et al.* A hexanucleotide repeat expansion in C9ORF72 is the cause of chromosome 9p21-linked ALS-FTD. *Neuron***72,** 257–268 (2011).

11. Beck, J. *et al.* Large C9orf72 hexanucleotide repeat expansions are seen in multiple neurodegenerative syndromes and are more frequent than expected in the UK population. *Am. J. Hum. Genet.* **92,** 345–353 (2013).
12. van Blitterswijk, M. *et al.* Association between repeat sizes and clinical and pathological characteristics in carriers of C9ORF72 repeat expansions (Xpansize-72): a cross-sectional cohort study. *Lancet Neurol* **12,** 978–988 (2013).
13. Šket, P. *et al.* Characterization of DNA G-quadruplex species forming from C9ORF72 G4C2-expanded repeats associated with amyotrophic lateral sclerosis and frontotemporal lobar degeneration. *Neurobiology of Aging* **36,** 1091–1096 (2015).
14. Xi, Z. *et al.* The C9orf72 repeat expansion itself is methylated in ALS and FTLD patients. *Acta Neuropathol.* **129,** 715–727 (2015).
15. Verkerk, A. J. *et al.* Identification of a gene (FMR-1) containing a CGG repeat coincident with a breakpoint cluster region exhibiting length variation in fragile X syndrome. *Cell* **65,** 905–914 (1991).
16. Paulson, H. Repeat expansion diseases. *Handbook of Clinical Neurology* **147,** 105–123 (2018).
17. Van Mossevelde, S. *et al.* Clinical Evidence of Disease Anticipation in Families Segregating a C9orf72 Repeat Expansion. *JAMA Neurol* **74,** 445–452 (2017).
18. Song, Q. *et al.* A Reference Methylome Database and Analysis Pipeline to Facilitate Integrative and Comparative Epigenomics. *PLoS ONE* **8,** e81148 (2013).
19. Xi, Z. *et al.* Hypermethylation of the CpG Island Near the G4C2 Repeat in ALS with a C9orf72 Expansion. *The American Journal of Human Genetics* **92,** 981–989 (2013).
20. Russ, J. *et al.* Hypermethylation of repeat expanded C9orf72 is a clinical and molecular disease modifier. *Acta Neuropathol.* **129,** 39–52 (2015).
21. Cohen-Hadad, Y. *et al.* Marked Differences in C9orf72 Methylation Status and Isoform Expression between C9/ALS Human Embryonic and Induced Pluripotent Stem Cells. *Stem Cell Reports* **0,** (2016).
22. Esanov, R. *et al.* A C9ORF72 BAC mouse model recapitulates key epigenetic perturbations of ALS/FTD. *Mol Neurodegeneration* **12,** 46 (2017).
23. Bell, M. V. *et al.* Physical mapping across the fragile X: Hypermethylation and clinical expression of the fragile X syndrome. *Cell* **64,** 861–866 (1991).
24. Hornstra, L. K., Nelson, D. L., Warren, S. T. & Yang, T. P. High resolution methylation analysis of the FMR1 gene trinucleotide repeat region in fragile X syndrome. *Hum Mol Genet* **2,** 1659–1665 (1993).
25. Xi, Z. *et al.* Hypermethylation of the CpG-island near the C9orf72 $G_4C_2$-repeat expansion in FTLD patients. *Hum Mol Genet* **23,** 5630–5637 (2014).
26. Monaghan, K. G., Lyon, E. & Spector, E. B. ACMG Standards and Guidelines for fragile X testing: a revision to the disease-specific supplements to the Standards and Guidelines for Clinical Genetics Laboratories of the American College of Medical Genetics and Genomics. *Genetics in Medicine 2013 15:7* **15,** 575–586 (2013).
27. Lyons, J. I., Kerr, G. R. & Mueller, P. W. Fragile X Syndrome: Scientific Background and Screening Technologies. *The Journal of Molecular Diagnostics* **17,** 463–471 (2015).
28. Cleary, E. M. *et al.* Improved PCR based methods for detecting C9orf72 hexanucleotide repeat expansions. *Molecular and Cellular Probes* **30,** 218–224 (2016).
29. Bahlo, M. *et al.* Recent advances in the detection of repeat expansions with short-read next-generation sequencing. *F1000Res* **7,** (2018).
30. Almeida, S. *et al.* Modeling key pathological features of frontotemporal dementia with C9ORF72 repeat expansion in iPSC-derived human neurons. *Acta Neuropathol.* **126,** 385–399 (2013).
31. McFarland, K. N. *et al.* SMRT Sequencing of Long Tandem Nucleotide Repeats in SCA10 Reveals Unique Insight of Repeat Expansion Structure. *PLoS ONE* **10,** e0135906 (2015).

32. Ebbert, M. T. W. *et al.* Long-read sequencing across the C9orf72 'GGGGCC' repeat expansion: implications for clinical use and genetic discovery efforts in human disease. *Mol Neurodegeneration* **13,** 46 (2018).
33. Schüle, B. *et al.* Parkinson's disease associated with pure ATXN10 repeat expansion. *npj Parkinson's Disease 2017 3:1* **3,** 27 (2017).
34. Ishiura, H. *et al.* Expansions of intronic TTTCA and TTTTA repeats in benign adult familial myoclonic epilepsy. *Nat Genet* **92,** 1 (2018).
35. Tsai, Y.-C. *et al.* Amplification-free, CRISPR-Cas9 Targeted Enrichment and SMRT Sequencing of Repeat-Expansion Disease Causative Genomic Regions. *bioRxiv* 203919 (2017). doi:10.1101/203919
36. Mertes, F. *et al.* Targeted enrichment of genomic DNA regions for next-generation sequencing. *Brief Funct Genomics* **10,** 374–386 (2011).
37. Simpson, J. T. *et al.* Detecting DNA cytosine methylation using nanopore sequencing. *Nat Methods* (2017). doi:10.1038/nmeth.4184
38. Schatz, M. C. Nanopore sequencing meets epigenetics. *Nat Methods* **14,** 347–348 (2017).
39. Rand, A. C. *et al.* Mapping DNA methylation with high-throughput nanopore sequencing. *NatMethods* (2017). doi:10.1038/nmeth.4189
